# ORAI1 establishes resistance to SARS-CoV-2 infection by regulating tonic type I interferon signaling

**DOI:** 10.1101/2021.05.04.442548

**Authors:** Beibei Wu, Arunachalam Ramaiah, Gustavo Garcia, Yousang Gwack, Vaithilingaraja Arumugaswami, Sonal Srikanth

**Author notes:** Senior authors (Vaithilingaraja Arumugaswami, Sonal Srikanth). Equal contribution (Beibei Wu, Arunachalam Ramaiah). Corresponding authors, Address correspondence to: Dr. Sonal Srikanth, Department of Physiology, David Geffen School of Medicine, 53-266 CHS, 10833 Le Conte Avenue, Los Angeles, CA 90095, or Dr. Vaithilingaraja Arumugaswami, B2-049A CHS, Box 956948, University of California, Los Angeles, CA 90095.

## Abstract

ORAI1 and STIM1 are the critical mediators of store-operated Ca^2+^ entry by acting as the pore subunit and an endoplasmic reticulum-resident signaling molecule, respectively. In addition to Ca^2+^ signaling, STIM1 is also involved in regulation of a cytosolic nucleic acid sensing pathway. Using *ORAI1* and *STIM1* knockout cells, we examined their contribution to the host response to SARS-CoV-2 infection. *STIM1* knockout cells showed strong resistance to SARS-CoV-2 infection due to enhanced type I interferon response. On the contrary, *ORAI1* knockout cells showed high susceptibility to SARS-CoV-2 infection as judged by increased expression of viral proteins and a high viral load. Mechanistically, *ORAI1* knockout cells showed reduced homeostatic cytoplasmic Ca^2+^ concentration and severe impairment in tonic interferon signaling. Transcriptome analysis showed downregulation of multiple cellular defense mechanisms, including antiviral signaling pathways in ORAI1 knockout cells, which are likely due to reduced expression of the Ca^2+^-dependent transcription factors of the activator protein 1 (AP-1) family and *MEF2C*. Our results identify a novel role of ORAI1-mediated Ca^2+^ signaling in regulating the baseline type I interferon level, which is a determinant of host resistance to SARS-CoV-2 infection.

## Introduction

Type I Interferon (IFN-I) response provides the first line of defense against viral infection (tenOever, 2016), and is also important for the development of adaptive immunity (Iwasaki and Medzhitov, 2015). IFN-I response is ubiquitous in almost all nucleated cells, whereas type III IFNs (IFN-III) are restricted to anatomic barriers and specific immune cells (Lazear et al., 2019). This selectivity is due to the receptor expression patterns: IFN-I binds IFN-α receptor (IFNAR) 1 and IFNAR2, which are ubiquitously expressed, while type III IFN binds IFNLR1 and IL-10-Rβ, which are preferentially expressed in epithelial cells (Lazear et al., 2019). Despite using different receptors, IFN-I and IFN-III activate the same downstream signaling pathway and induce transcription of hundreds of IFN-stimulated genes (ISGs), with IFN-III inducing lower levels of ISG expression than IFN-I (Crotta et al., 2013). The importance of IFN signaling pathway is underscored by the fact that most viruses have developed mechanisms to inactivate this pathway (Schulz and Mossman, 2016).

Coronavirus disease 2019 (COVID-19) pandemic is one of the worst crises of our times, prompting an urgent need to uncover host defense mechanisms to its causative agent, severe acute respiratory syndrome coronavirus 2 (SARS-CoV-2). SARS-CoV-2 utilizes multiple approaches to evade host IFN response, including suppression of IFN-I production as well as IFN-I signaling (Xia and Shi, 2020). The clinical manifestations and pathology of COVID-19 are similar to those of SARS-CoV-1, but the severities of the diseases differ (Xie and Chen, 2020). Unlike severe SARS-CoV-1 infection, SARS-CoV-2 infection shows a wide range of clinical features, ranging from asymptomatic, mild and moderate to severe and critical. The clinical manifestations of virus infection depend on virus-host interactions. Asymptomatic outcomes of SARS-CoV-2 infection may thus be attributed to strong host innate antiviral defense. In clinics, COVID-19 patients with circulating antibodies to IFNI or loss-of-function mutations in genes necessary to mount the IFN-I response developed life-threatening pneumonia (Bastard et al., 2020; Zhang et al., 2020). Recent studies have found that unlike original coronaviruses, SARS-CoV-2 is highly sensitive to pre-treatment with type I and III IFNs. Pre-treatment of immortalized or primary human airway epithelial cells with IFN-I or IFN-III imparts resistance to SARS-CoV-2 infection (Blanco-Melo et al., 2020; Lokugamage et al., 2020; Vanderheiden et al., 2020). Taken together, a picture emerges that the baseline levels of IFN-I that is known to prime the host antiviral status (Gough et al., 2012) can also provide a significant barrier to SARS-CoV-2 infection.

Store-operated Ca^2+^ entry (SOCE), induced by depletion of the endoplasmic reticulum (ER) Ca^2+^ stores after activation of G protein-coupled receptors or receptor tyrosine kinases, is a ubiquitous mechanism of elevation of intracellular Ca^2+^ concentration ([Ca^2+^]) in most cell types. Ca^2+^ release-activated Ca^2+^ (CRAC) channels are the specialized class of store-operated Ca^2+^ (SOC) channels that play a primary role in the elevation of [Ca^2+^] in immune cells (Prakriya and Lewis, 2015; Srikanth and Gwack, 2013). CRAC channels consist of two major components, the plasma membrane (PM)-localized pore subunit, ORAI1, and an endoplasmic reticulum (ER)-resident Ca^2+^ sensor, stromal interaction molecule 1 (STIM1). STIM1 senses depletion of the ER Ca^2+^ stores and interacts with ORAI1 to open the pore. Ca^2+^ signaling mediated by CRAC channels is essential for induction of transcriptional programs via the NFAT (nuclear factor of activated T cells) pathway (Prakriya and Lewis, 2015; Srikanth and Gwack, 2013). Severe combined immune deficiency (SCID) caused by mutations in *ORAI1* or *STIM1*, and the widespread clinical use of inhibitors of this pathway - cyclosporine A (CsA) and FK506 as immunosuppressants, underscore the importance of therapeutic targeting of the Ca^2+^-NFAT pathway (Prakriya and Lewis, 2015; Srikanth and Gwack, 2013).

While the current understanding of the role of ORAI1 is limited to Ca^2+^ signaling, a Ca^2+^ signaling-independent role of STIM1 in regulating the basal IFN-I response was uncovered recently (Srikanth et al., 2019). Myeloid cell-specific *Stim1* knockout (KO) mice and a patient lacking STIM1 expression were shown to have elevated baseline serum IFN-β and increased expression of various ISGs. Accordingly, myeloid-specific *Stim1* KO mice showed enhanced resistance to DNA virus infection, which was not observed in *Orai1* KO cells and mice (Srikanth et al., 2019). Here, we examined the role of *ORAI1* and *STIM1* in host resistance to infection with an RNA virus, SARS-CoV-2. Loss of ORAI1 markedly reduced the host resistance to SARS-CoV-2 by abolishing the baseline IFN-β levels and impairing tonic IFN signaling. On the contrary, *STIM1* KO cells showed remarkable resistance to SARS-CoV-2 infection supporting previous studies of enhanced baseline IFN-I signaling (Srikanth et al., 2019). Our data suggest ORAI1-mediated Ca^2+^ signaling is crucial for tonic IFN-I signaling, which primes the cellular antiviral state and thereby determines host resistance to SARS-CoV-2 infection.

## Results

### Loss of ORAI1 reduces store-operated Ca^2+^ entry and the baseline cytoplasmic Ca^2+^ concentration in HEK293-ACE2 cells

To examine the role of ORAI1 and STIM1 in cellular response to SARS-CoV-2, we generated HEK293 cells stably expressing the receptor for SARS-CoV-2, angiotensin converting enzyme 2 (ACE2) (Li et al., 2003). The resulting HEK293-ACE2 cells were transduced with lentiviruses encoding Cas9 and sgRNAs targeting either *ORAI1* or *STIM1*. All three lines expressed similar levels of ACE2 as judged by immunocytochemistry and immunoblotting (**Suppl. Figs. 1A and 1B**). Loss of STIM1 protein expression was validated by immunoblotting, while that of ORAI1 was validated by flow cytometry (**Figs. 1A and 1B**). Multiple studies have shown loss of SOCE in HEK293 cells deleted for expression of *ORAI1/STIM1* (Nomura et al., 2020; Yoast et al., 2020). To measure SOCE in *ORAI1*^-/-^ and *STIM1*^-/-^ HEK293-ACE2 cells, we used thapsigargin, a blocker of SERCA (sarco/endoplasmic reticulum Ca^2+^-ATPase) pump, that allows for passive depletion of the ER Ca^2+^ stores, thereby activating SOCE. We observed almost complete abrogation of SOCE in both *ORAI1*^-/-^ and *STIM1*^-/-^ HEK293-ACE2 cells (**Fig. 1C**). While control cells showed a uniform increase in SOCE after re-introducing Ca^2+^-containing Ringer’s solution, more than 95% of the polyclonal *ORAI1*^-/-^ and *STIM1*^-/-^ HEK293-ACE2 cells remained unresponsive. The Ca^2+^ release from the ER in those cells was similar. Next, we checked whether loss of ORAI1/STIM1 affected resting cytoplasmic [Ca^2+^] in unstimulated cells. Using our previously established protocols, we sequentially perfused the cells with Ca^2+^-free, 2 mM, and 20 mM Ca^2+^-containing Ringer’s solutions without any store depletion, to measure the homeostatic Ca^2+^ levels in cells (Srikanth et al., 2010a). Interestingly, in these measurements, only *ORAI1*^-/-^ HEK293-ACE2 cells showed significant reduction in baseline [Ca^2+^] in the presence of 2- or 20-mM Ca^2+^-containing Ringer’s solution, suggesting an additional role of ORAI1 in regulating basal cytoplasmic [Ca^2+^] in these cells (**Fig. 1D**). These data are in concurrence with previous observations where STIM2, a homolog of STIM1, was shown to be involved in maintaining resting cytoplasmic [Ca^2+^] together with ORAI1 (Brandman et al., 2007). In those experiments, STIM2-depleted cells showed lowered resting cytoplasmic [Ca^2+^], while those depleted for STIM1 had normal levels of resting cytoplasmic [Ca^2+^]. Collectively, by stable expression of ACE2 and deletion of *ORAI1/STIM1*, we developed a system to examine the role of ORAI1/STIM1 in the host response to SARS-CoV-2 infection and identified the role of ORAI1 in Ca^2+^ homeostasis in these cells.

**Figure 1.**
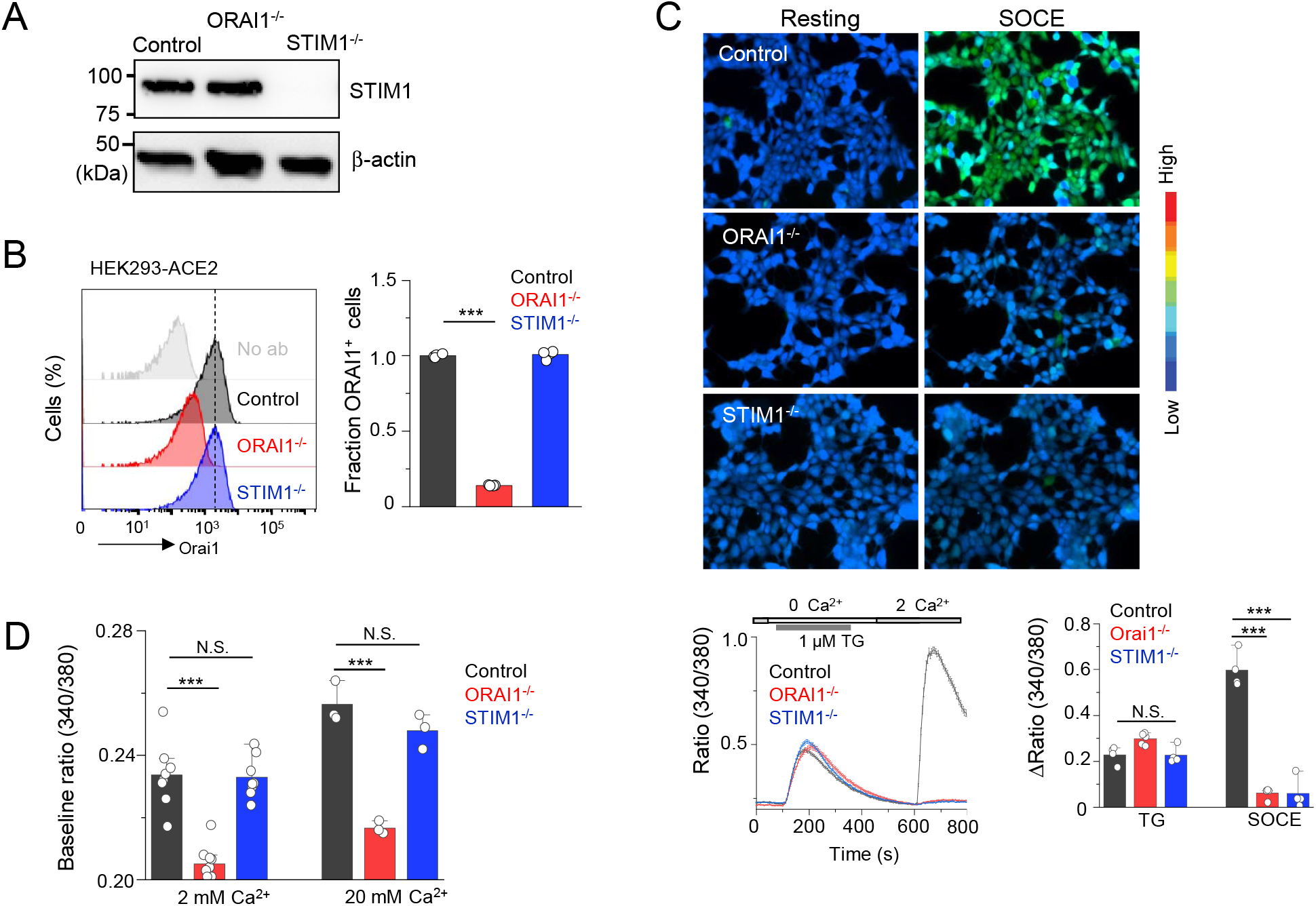
Loss of ORAI1 reduces basal Ca^2+^ concentration and abrogates SOCE in HEK293-ACE2 cells. (**A**) Representative immunoblot showing expression of STIM1 in lysates from control, *ORAI1*^-/-^, or *STIM1*^-/-^ HEK293-ACE2 cells. β-actin – loading control. Data are representative of two independent experiments. (**B**) Representative histograms showing levels of total ORAI1 protein in control, *ORAI1*^-/-^, and *STIM1*^-/-^ HEK293-ACE2 cells after permeabilization and intracellular staining with anti-ORAI1 antibody. The bar graph shows average (± s.e.m.) from three independent experiments. (**C**) Representative pseudocoloured epifluorescence images of indicated cells under resting conditions or at the peak of SOCE. Below - Representative traces showing averaged SOCE from control (39 cells), *ORAI1*^-/-^ (35 cells) and *STIM1*^-/-^ (42 cells) HEK293-ACE2 cells after passive depletion of intracellular Ca^2+^ stores with thapsigargin (TG – 1 μM) in Ca^2+^-free external solution. SOCE was measured after replacing external solution with that containing 2 mM CaCl_2_. Bar graph on the right shows averaged baseline subtracted ER Ca^2+^ stores (TG) and SOCE (± s.e.m.) from four independent experiments. (**D**) Bar graph shows baseline Ca^2+^ levels (as depicted by Fura-2 ratio) ± s.e.m. in indicated cell types upon perfusion with extracellular solution containing either 2 mM or 20 mM CaCl_2_. Each dot represents data obtained from an independent experiment. \*\*\**P*<0.0005 (two-tailed *t* test).

### Loss of ORAI1/STIM1 affects host resistance to SARS-CoV-2 infection

Next, we infected control, *ORAI1*^-/-^, and *STIM1*^-/-^ HEK293-ACE2 cells with SARS-CoV-2 at two different multiplicity of infections (MOIs) – 0.1 and 1.0. Twenty hours after infection, cells were harvested to examine expression of viral proteins by immunoblotting and immunocytochemistry, as well as quantification of viral genome. We observed high levels of viral protein expression in SARS-CoV-2-infected *ORAI1*^-/-^ HEK293-ACE2 cells when compared to control cells by immunoblotting (**Fig. 2A**), such that twenty hours after infection at high MOI (1.0), a significant fraction of the *ORAI1*^-/-^ cells were lysed (**Fig. 2B**). On the contrary, we observed strong resistance of *STIM1*^-/-^ HEK293-ACE2 cells to SARS-CoV-2 infection. *STIM1*^-/-^ HEK293-ACE2 cells showed markedly reduced expression of viral protein both by immunoblotting and immunocytochemistry when compared to control cells (**Figs. 2A and 2B**). Accordingly, we observed fewer plaques in *STIM1*^-/-^ cells when compared to control cells. Genome copy measurements at five hours after infection showed significantly increased and decreased susceptibility of *ORAI1*^-/-^ and *STIM1*^-/-^ cells, respectively, especially in cells infected at high MOI (**Fig. 2C**, left). Further, the increased and decreased susceptibility of *ORAI1*^-/-^ and *STIM1*^-/-^ cells, respectively, was obvious at twenty hours after infection, where *ORAI1*^-/-^ cells showed ~100-fold increase in viral genome copy numbers whereas *STIM1*^-/-^ cells showed more than 100-fold reduction, when compared to control HEK293-ACE2 cells (**Fig. 2C**, right).

**Figure 2.**
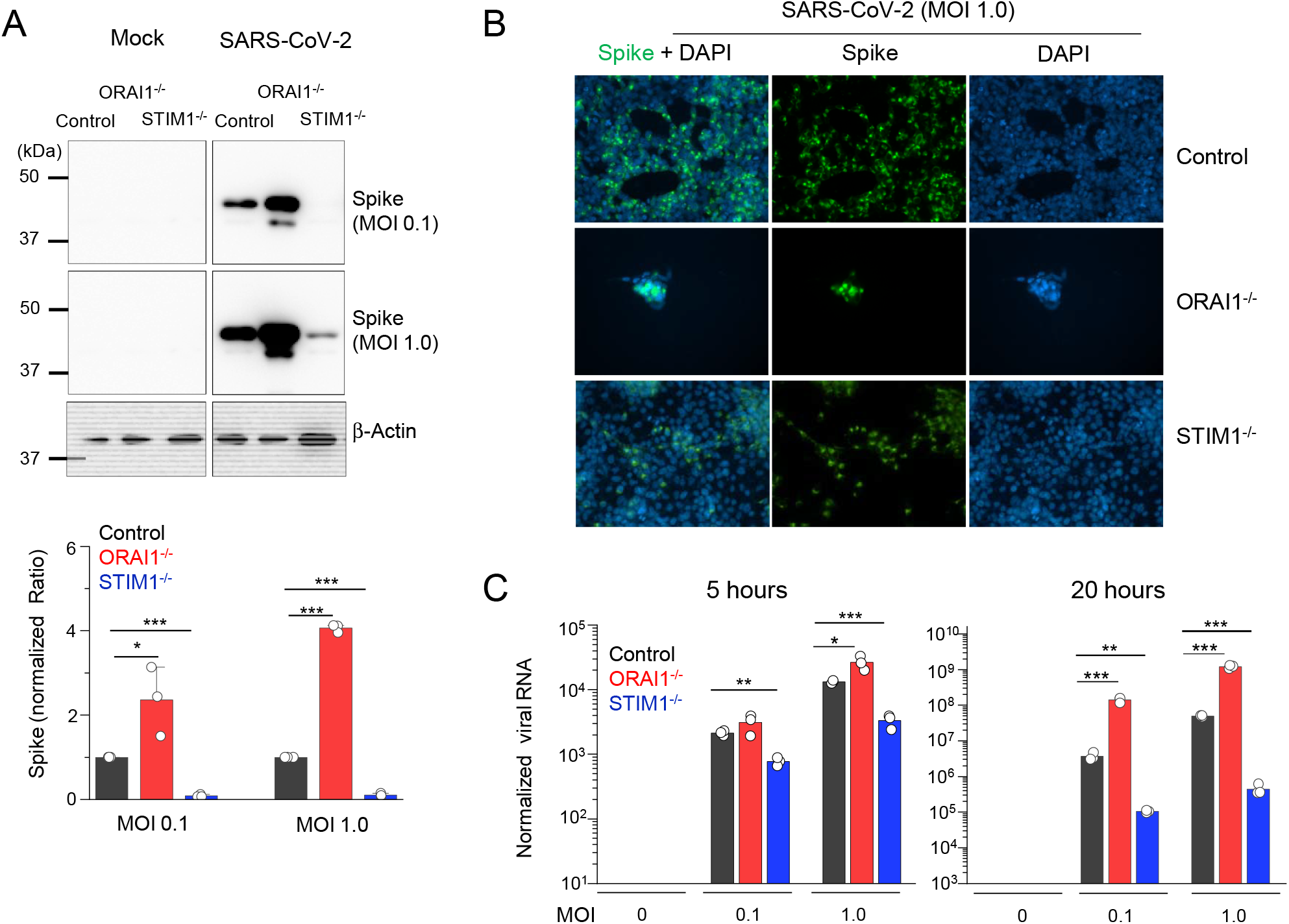
Loss of ORAI1 or STIM1 affects cellular susceptibility to SARS-CoV-2 infection. (**A**) Representative immunoblot showing expression of SARS-CoV-2 proteins in indicated HEK293-ACE2 cells under mock conditions or after infection with SARS-CoV-2 at indicated multiplicity of infection (MOIs). β-actin – loading control. Bar graph below shows densitometry analysis of normalized ratio of SARS-N-CoV-2 to β-actin (± s.e.m.) from three independent experiments. (**B**) Representative epifluorescence images showing expression of spike protein (green) in control, *ORAI1*^-/-^, or *STIM1*^-/-^ HEK293-ACE2 cells after infection with SARS-CoV-2 at MOI 1.0. Cells were co-stained with DAPI for detection of nuclei. *ORAI1*^-/-^ HEK293-ACE2 cells were all detached due to high virus load. (**C**) Quantitative RT-PCR analysis of viral genome from indicated cell types under mock conditions or after infection with SARS-CoV-2 at indicated MOI for 5 (left graph) or 20 (right graph) hours. Shown is one representative triplicate from two independent experiments. * *P*<0.05, ** *P*<0.005, \*\*\**P*<0.0005 (two-tailed *t* test).

### *STIM1* KO cells exhibit resistance to SARS-CoV-2 infection due to higher baseline type I IFN response

Based on our previous data showing pre-activation of the IFN-I pathway in *Stim1*^-/-^ fibroblasts and macrophages (Srikanth et al., 2019), and high sensitivity of SARS-CoV-2 to IFN-I (Blanco-Melo et al., 2020; Lokugamage et al., 2020; Vanderheiden et al., 2020), we surmised that the resistance of *STIM1*^-/-^ cells to SARS-CoV-2 infection was most likely derived from an increase in the baseline IFN-I response. As expected, *STIM1*^-/-^ HEK293-ACE2 cells indeed showed higher levels of baseline IFN-β levels, which remained elevated even after SARS-CoV-2 infection (**Fig. 3A**). We also observed enhanced IL-6 production by *STIM1*^-/-^ cells under resting conditions and after SARS-CoV-2 infection when compared to control cells (**Fig. 3B**). Notably, these results showed that SARS-CoV-2 infection did not trigger a robust IFN-I response but induced strong IL-6 expression in consistence with previous observations (Chen et al., 2020).

**Figure 3.**
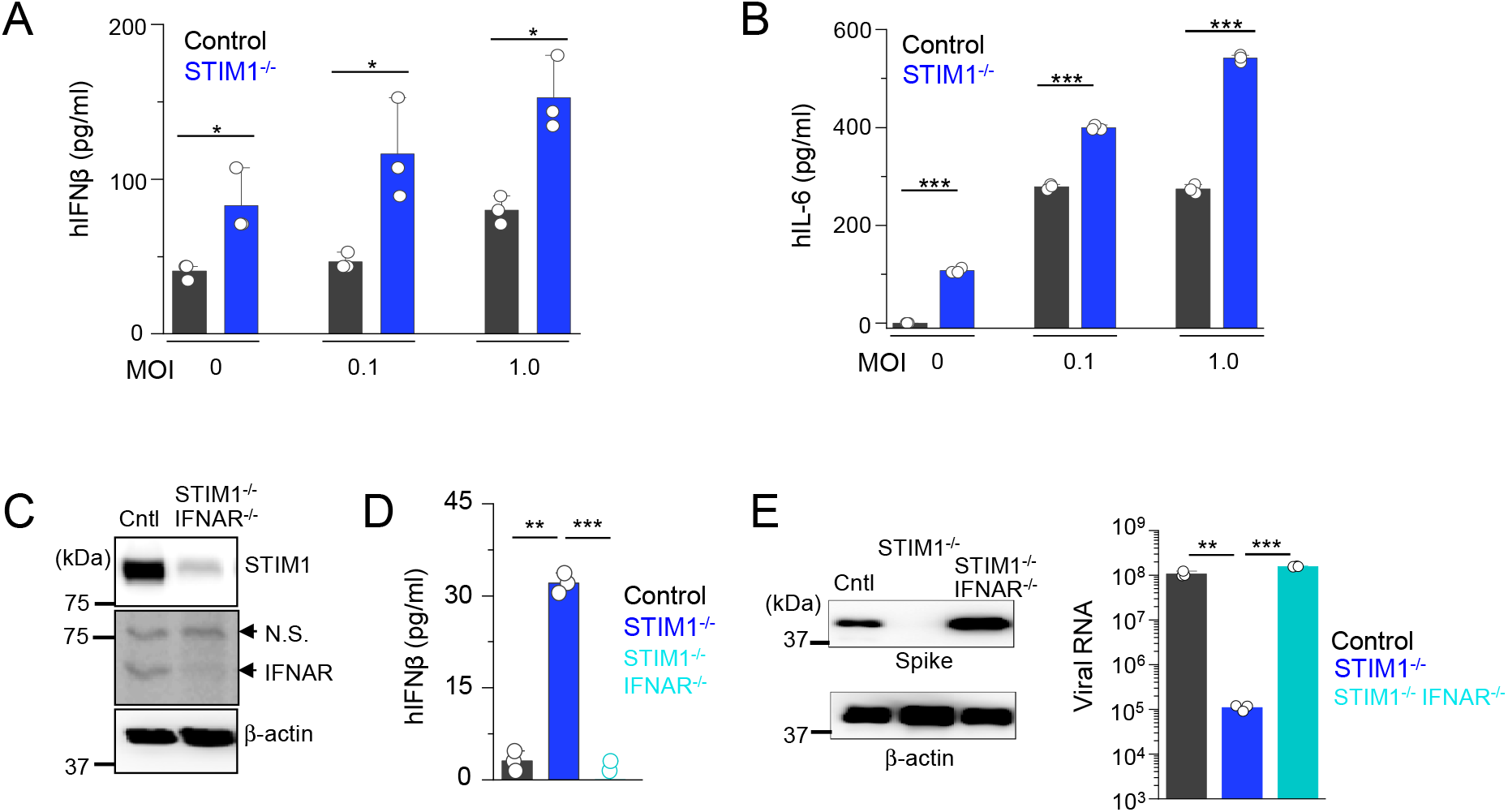
Loss of STIM1 imparts resistance to SARS-CoV-2 infection by enhancing interferon β signaling pathway. (**A**) Levels of IFN-β in culture supernatants from control or *STIM1*^-/-^, HEK293-ACE2 cells under mock conditions or 20 h after infection with SARS-CoV-2 at indicted MOIs. (**B**) Levels of IL-6 in culture supernatants from control or *STIM1*^-/-^ HEK293-ACE2 cells. (**C**) Representative immunoblot showing expression of STIM1 and interferon α receptor subunit 1 (IFNAR1) in control HEK293-ACE2 cells or those deleted for both STIM1 and IFNAR1 using CRISPR/Cas9-mediated recombination. β-actin – loading control. Data are representative of two independent experiments. N.S., non-specific band. (**D**) Levels of IFN-β in culture supernatants from control, *STIM1*^-/-^ or *STIM1*^-/-^ IFNAR^-/-^ HEK293-ACE2 cells under resting conditions. (**E**) Representative immunoblot showing expression of SARS-CoV-2 proteins in indicated HEK293-ACE2 cells under mock conditions or after infection with SARS-CoV-2 at MOI 0.1. β-actin – loading control. Data are representative of two independent infection experiments. Right - Quantitative RT-PCR analysis of viral genome from indicated cell types under mock conditions or after infection with SARS-CoV-2 at MOI 0.1 for 20 hours. In bar graphs in panels (**A**), (**B**), (**D**), and (**E**) representative triplicates from two independent experiments are shown. * *P*<0.05, ** *P*<0.005, \*\*\**P*<0.0005 (two-tailed *t* test).

To validate whether the resistance of *STIM1*^-/-^ cells to SARS-CoV-2 infection was due to elevated IFN-I signaling pathway, we generated HEK293-ACE2 cells deleted of both *STIM1* and *IFNAR1* using the CRISPR/Cas9 system (**Fig. 3C**). Deletion of *IFNAR1* abolished the increased baseline IFN-ß expression in *STIM1*^-/-^ cells (**Fig. 3D**). In addition, co-deletion of *IFNAR1* abolished the resistance of *STIM1*^-/-^ cells to SARS-CoV-2, as demonstrated by enhanced expression of viral proteins and increased burden of viral genome copies (**Fig. 3E**). These results suggest that the strong resistance of *STIM1*^-/-^ cells to SARS-CoV-2 is predominantly derived from enhanced basal level of IFN-β. Considering that SARS-CoV-2 did not trigger robust IFN-I signaling, these results also emphasize the importance of the basal IFN-I levels in host resistance to SARS-CoV-2 infection.

### High susceptibility of *ORAI1* KO cells to SARS-CoV-2 infection is due to low baseline type I IFN signaling

The increased susceptibility of *ORAI1*^-/-^ cells to SARS-CoV-2 infection was surprising because ORAI1 deficiency did not influence host resistance to infection with a DNA virus, herpes simplex virus type 1 (HSV-1) in a previous study (Srikanth et al., 2019). Because SARS-CoV-2 was highly sensitive to the baseline IFN-I levels as observed in our analysis of *STIM1*^-/-^ cells, we hypothesized that ORAI1 may influence the basal IFN-I levels. Culture supernatant analysis showed almost complete abrogation of baseline IFN-β levels in *ORAI1*^-/-^ cells when compared to control cells under resting conditions as well as upon infection with SARS-CoV-2 at low MOI (**Fig. 4A**). IL-6 levels in these cells were not influenced. In support of reduced IFN-β levels, *ORAI1*^-/-^ cells also showed reduced activation of the IFN-I signaling pathway, including reduced basal levels of p-STAT1 as well as a decrease in total STAT1 proteins (**Fig. 4B**), which has been proposed to be dependent on the baseline IFN-β signaling (Gough et al., 2012).

**Figure 4.**
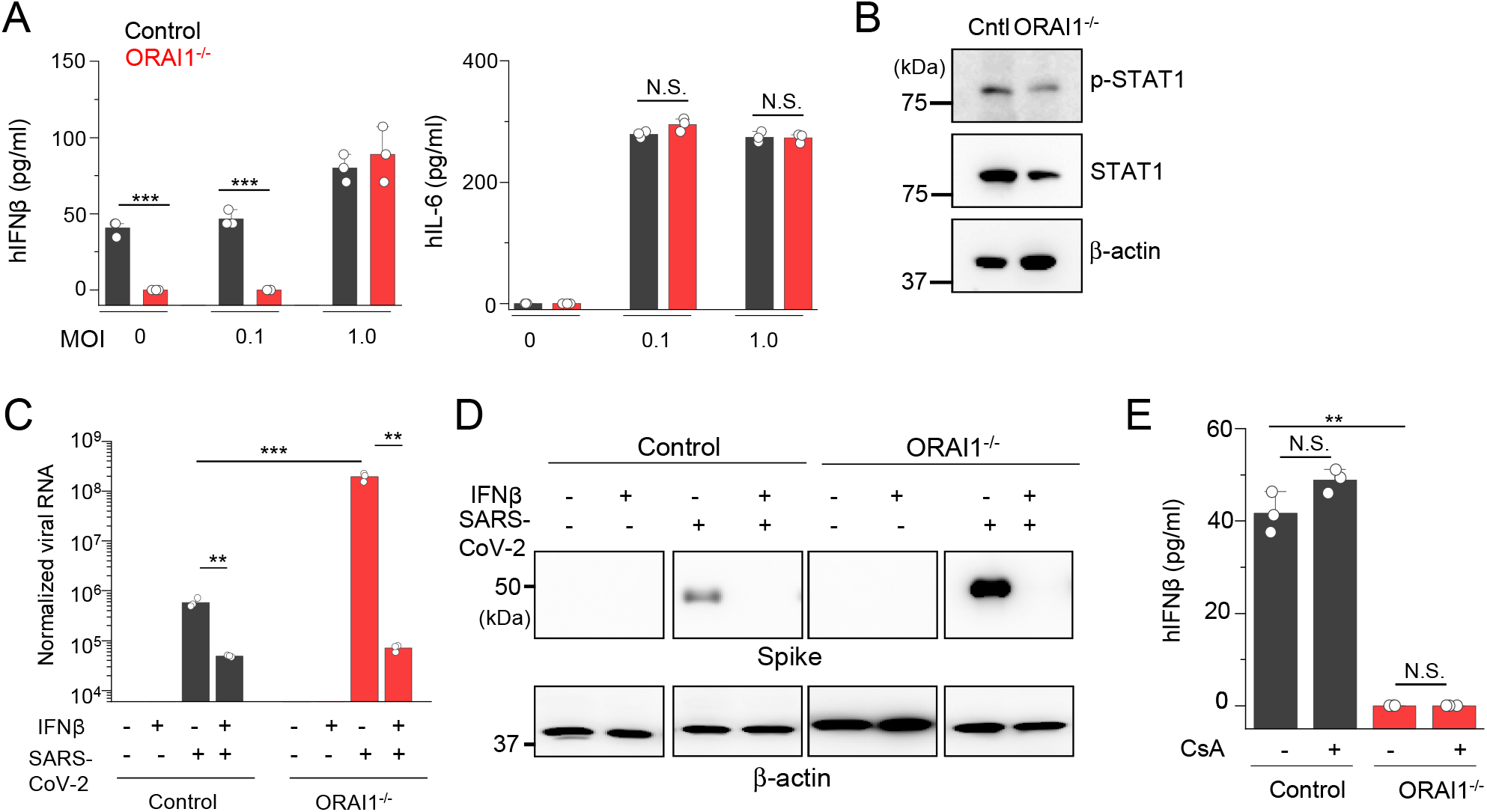
Loss of ORAI1 imparts susceptibility to SARS-CoV-2 infection by abrogating baseline interferon β levels. (**A**) Levels of IFNβ and IL-6 in culture supernatants from control or *ORAI1*^-/-^ HEK293-ACE2 cells under mock conditions or 20 hours after infection with SARS-CoV-2 at indicted MOIs. (**B**) Representative immunoblot showing expression of phosphorylated STAT1 (p-STAT1) or total STAT1 in lysates from control or *ORAI1*^-/-^ HEK293-ACE2 cells. β-actin – loading control. Data are representative of three independent experiments. (**C**) Quantitative RT-PCR analysis of viral genome from indicated cell types with or without pre-treatment with 100 U/ml of IFN-β (for 20 hours) under mock conditions or after infection with SARS-CoV-2 at MOI 0.1 for 20 h. (**D**) Representative immunoblot showing expression of SARS-CoV-2 proteins in indicated HEK293-ACE2 cells with or without pre-treatment with 100 U/ml of IFN-β (for 20 hours) under mock conditions or after infection with SARS-CoV-2 at MOI 0.1. β-actin – loading control. Data are representative of two independent infection experiments. (**E**) Levels of IFN-β in culture supernatants from control or *ORAI1*^-/-^ HEK293-ACE2 cells with or without pretreatment with 1μM cyclosporine A (CsA, 20 hours). *P*<0.05, ** *P*<0.005, \*\*\**P*<0.0005 (two-tailed *t* test).

To check whether the high susceptibility of *ORAI1*^-/-^ cells to SARS-CoV-2 infection was rescued by treatment with IFN-β, we pre-treated these cells with low levels of IFN-β for ~20 hours before SARS-CoV-2 infection. IFN-β pre-treatment reduced the viral genome by ~10-fold in control cells, in agreement with other studies (Blanco-Melo et al., 2020; Lokugamage et al., 2020; Vanderheiden et al., 2020) (**Fig. 4C**). Importantly, pre-treatment of *ORAI1*^-/-^ cells with IFN-β made the cells highly resistant, with more than 1,000-fold reduction in viral genome copy number, similar to the levels observed with control cells. These observations were validated by immunoblotting to check expression of viral proteins, where pre-treatment with IFN-β abrogated expression of viral proteins equally in control and *ORAI1*^-/-^ cells (**Fig. 4D**). Since IFN-β pre-treatment in *ORAI1*^-/-^ cells almost completely rescued the phenotypes to the level in control cells, these data suggest that reduced IFN-I baseline may have a significant contribution towards high susceptibility of *ORAI1*^-/-^ cells to SARS-CoV-2 infection. Since the NFAT family of transcription factors are well-known downstream effectors of ORAI1-mediated SOCE, we checked whether reduced IFN-β levels in *ORAI1*^-/-^ cells were due to loss of NFAT function. To check this possibility, we used CsA, which blocks calcineurin, a phosphatase necessary for de-phosphorylation of NFAT in the cytoplasm. We pretreated control and *ORAI1*^-/-^ HEK293-ACE2 cells with CsA and examined its effect on the basal IFN-β production. Surprisingly, CsA treatment did not affect IFN-β levels in either of these cell types, suggesting a potential role of Ca^2+^-dependent transcription factors other than NFAT in this event (**Fig. 4E**).

### ORAI1 acts on Ca^2+^-regulated transcription factors to modulate the baseline IFN-I production

To uncover the antiviral defense mechanisms impaired in *ORAI1*^-/-^ cells, we performed transcriptome analysis by bulk RNA sequencing of control, *ORAI1*^-/-^, and *STIM1*^-/-^ HEK293-ACE2 cells before and 20 hours after SARS-CoV-2 infection (**Suppl. Fig. 2**). Among infected samples, viral count analysis showed uniform increase in expression of all the viral genes in *ORAI1*^-/-^ cells, while the same was uniformly reduced in *STIM1*^-/-^ cells, when compared to control cells (**Fig. 5A**). Importantly, the viral genome reads comprised ~20% of the total reads for control cells, ~77% for *ORAI1*^-/-^ cells, and <1% for *STIM1*^-/-^ cells, in concurrence with the observed phenotypes (**Fig. 5B**). Pathway enrichment analysis of *ORAI1*^-/-^ cells before infection showed downregulation of multiple antiviral defense mechanisms, including ISG15 antiviral mechanism and MyD88 signaling pathways (**Figs. 5C, D, and E**). Transcriptome analysis further identified multiple transcription factors known to be involved in antiviral pathways, whose expression was altered by loss of *ORAI1* (**Fig. 5F**). Among the transcription factors that were significantly downregulated in *ORAI1*^-/-^ cells, we identified *MEF2C*, and two members of the AP-1 family of transcription factors: *FOS*, and *ATF2*; all of which are known to be Ca^2+^-dependent for their functions (Ban et al., 2000; Lesch et al., 2015; McKinsey et al., 2002; Ng et al., 2009). The AP-1 family of transcription factors; including ATF2 and FOS are further known to bind the *IFNB* promoter to regulate its expression and may be involved in tonic IFN signaling (Gough et al., 2012). Remarkably, promoter analysis showed profound enrichment of MEF2C, FOS, and ATF2-bindings sites among the differentially expressed genes in *ORAI1*^-/-^ cells compared to the reference genome (**Fig. 5G**). Quantitative RT-PCR analysis confirmed downregulation of these three transcription factors in *ORAI1*^-/-^ cells under sterile conditions (**Fig. 5H**). Together, these data suggest that ORAI1 activity upregulates expression of the transcription factors *MEF2C*, *FOS* and *ATF2* leading to induction of multiple genes involved in antiviral defense mechanisms, including the baseline IFN-I signaling, providing resistance to SARS-CoV-2 infection.

**Figure 5.**
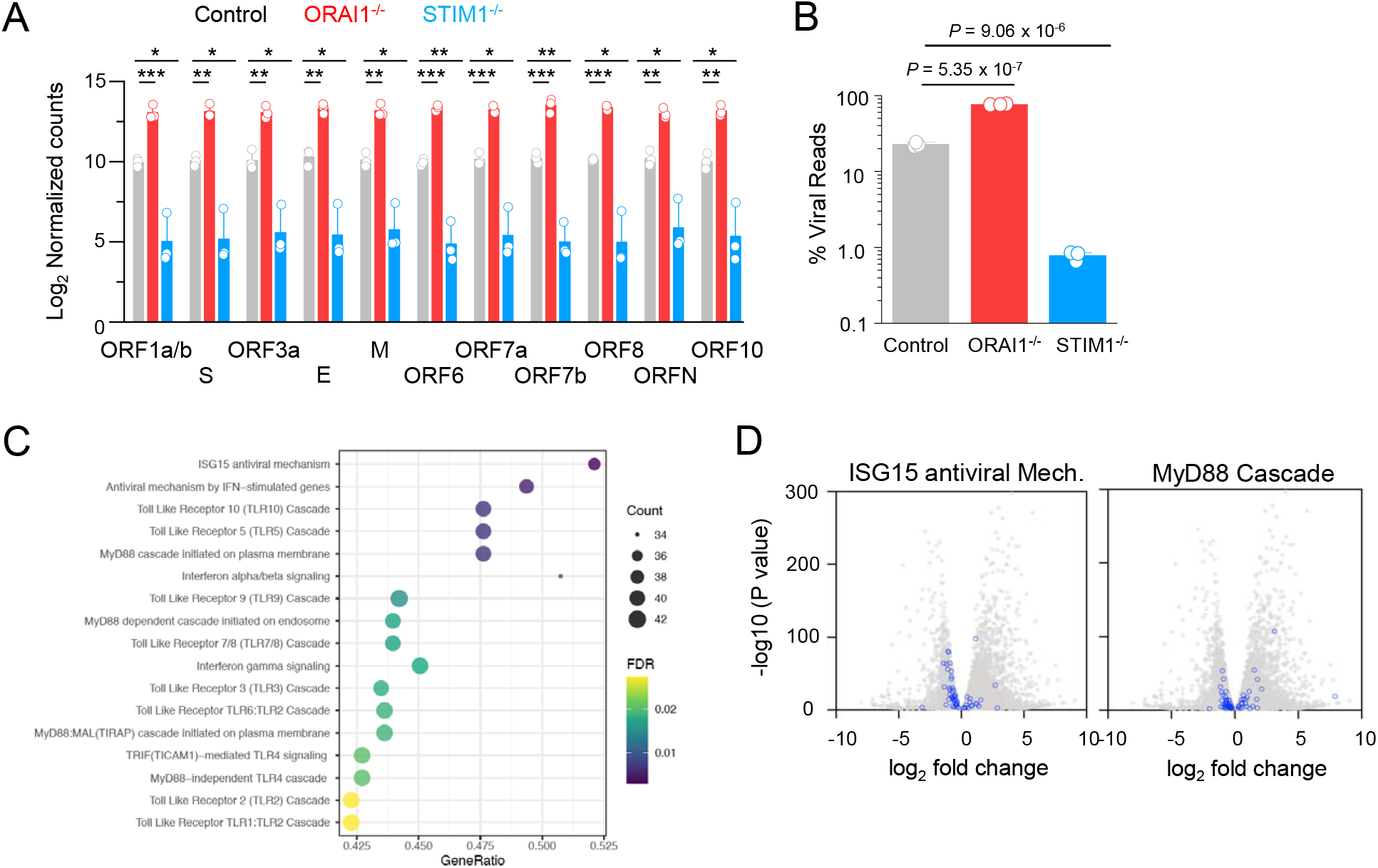
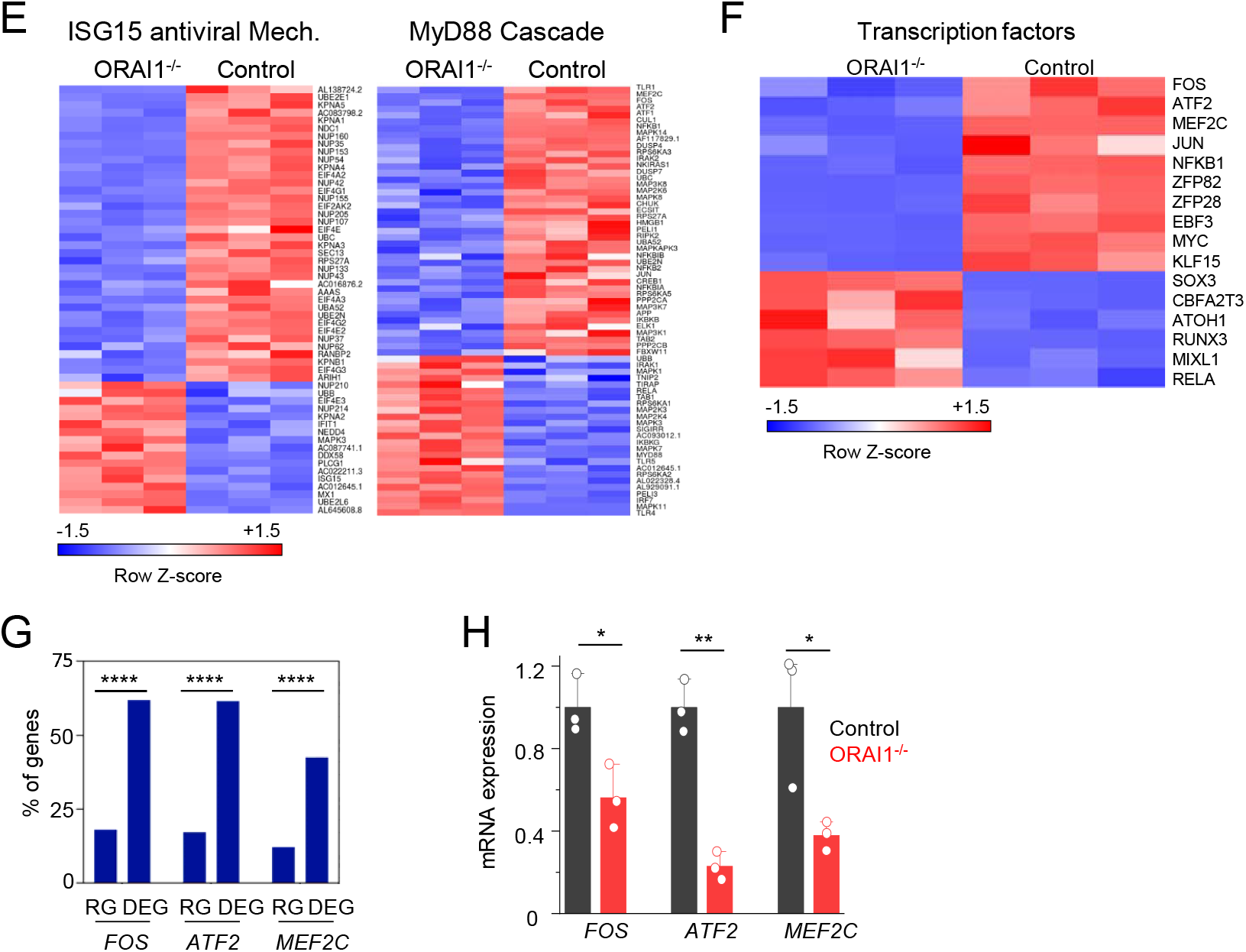
Transcriptome analysis of control and *ORAI1*^-/-^ HEK293-ACE2 cells. (**A**) Normalized read counts (log_2_) of SARS-CoV-2 RNA products, showing transcriptional enrichment of viral genes in SARS-CoV-2 infected cells when compared to uninfected cells. (**B**) Bar graph shows proportion of total reads comprising SARS-CoV-2 transcripts in indicated cell types. The proportion of virus-aligned reads over total reads is shown for each sample. Error bars represent average (± s.d.m.) from three biological replicates. (**C**) Dot plot visualization of enriched pathways in *ORAI1*^-/-^ HEK293-ACE2 cells. Reactome pathway enrichment analysis was performed in PANTHER. The size of the dots represents the percentage of genes enriched in the total gene set, while its color represents the false discovery rate (FDR, p-adjusted) value for each enriched pathway. (**D**) Volcano plots with all DEGs from *ORAI1*^-/-^ and control mock (uninfected) samples in gray and the indicated gene sets of anti-viral ISG15 signaling (left) and MyD88 signaling (right) highlighted in blue (FDR <0.01). (**E**) Heat map illustrating z-scores as expression levels of the differentially expressed genes involved in antiviral signaling and MyD88 signaling pathways as show in panel D. Blue and red colors represent down- and up-regulated genes, respectively. (**F**) Heat map depicting z-scores as expression levels of selected transcription factors differentially expressed between control and *ORAI1*^-/-^ HEK293-ACE2 cells. (**G**) Transcription factor binding sites at the promoters of genes (TSS ± 1 Kb) of the entire genome (from ChiP Atlas) or among differentially expressed genes (DEG) in *ORAI1*^-/-^ HEK293-ACE2 cells. The hypergeometric *P* values (**** P<0.0001) were calculated by comparing background (complete gene sets of the reference host; RG) to genes from DEGs to see enrichment of binding sites of indicated transcription factors. (**H**) Quantitative RT-PCR analysis of indicated genes from control and *ORAI1*^-/-^ HEK293-ACE2 cells under resting conditions. *P*<0.05, ** *P*<0.005, \*\*\**P*<0.0005 (two-tailed *t* test).

Recent studies using genome-wide CRISPR/cas9 knockout screens with SARS-CoV-2 infection have identified a plethora of host factors important for productive infection by SARS-CoV-2 (Daniloski et al., 2021; Hoffmann et al., 2021; Schneider et al., 2021; Wang et al., 2021; Wei et al., 2021). The list of necessary host factors required for SARS-CoV-2 propagation varied widely among the different screens, likely due to different cell types and infection conditions used for each screen. Not surprisingly, all the genome-wide screens identified ACE2, the receptor for SARS-CoV-2 as an essential factor for virus propagation. Our transcriptome analysis identified many of these host factors among the differentially expressed genes (DEGs) in *ORAI1*^-/-^ cells (**Suppl. Fig. 3**). However, many of these factors were downregulated in *ORAI1*^-/-^ cells, suggesting they may not play a significant role in high susceptibility to SARS-CoV-2 infection observed in these cells. Collectively, these results suggest that ORAI1-mediated Ca^2+^ signaling is mainly important for the regulation of the IFN-I levels, rather than host factors for SARS-CoV-2 infection.

### High sensitivity of SARS-CoV-2 to baseline IFN-I levels when compared to vesicular stomatitis virus

We next sought to examine whether ORAI1-mediated regulation of baseline IFN signaling played an important role in host response to a different RNA virus. For the same, we generated *ORAI1*^-/-^ and *STIM1*^-/-^ A549 cells (human lung epithelial cell-line) using the CRISPR/Cas9 system (**Figs. 6A and B**) and infected them with a negative-strand RNA virus, vesicular stomatitis virus (VSV)-GFP, at multiple different MOIs. Similar to our results with SARS-CoV-2, *ORAI1*^-/-^ A549 cells showed higher susceptibility to VSV-GFP infection, and *STIM1*^-/-^ A549 cells showed significant resistance at multiple different MOIs (**Fig. 6C**). However, *ORAI1*^-/-^ cells seem to be less sensitive to VSV-GFP infection when compared to SARS-CoV-2 infection. While *ORAI1*^-/-^ SARS-CoV-2-infected cells showed 3-4-fold higher expression of spike protein than control cells, the GFP^+^ population was at the most 1.5-2-fold higher than controls in *ORAI1*^-/-^ VSV-GFP-infected cells (Figs. **2A** and **6C**). These data confirm previous observations of exceptionally high sensitivity of SARS-CoV-2 to IFN-I treatment (Blanco-Melo et al., 2020; Lokugamage et al., 2020; Vanderheiden et al., 2020). Further, these data emphasize the significance of baseline IFN-I signaling-mediated priming in the host resistance to SARS-CoV-2 infection and the critical role of ORAI1-mediated Ca^2+^ signaling in this process.

**Figure 6.**
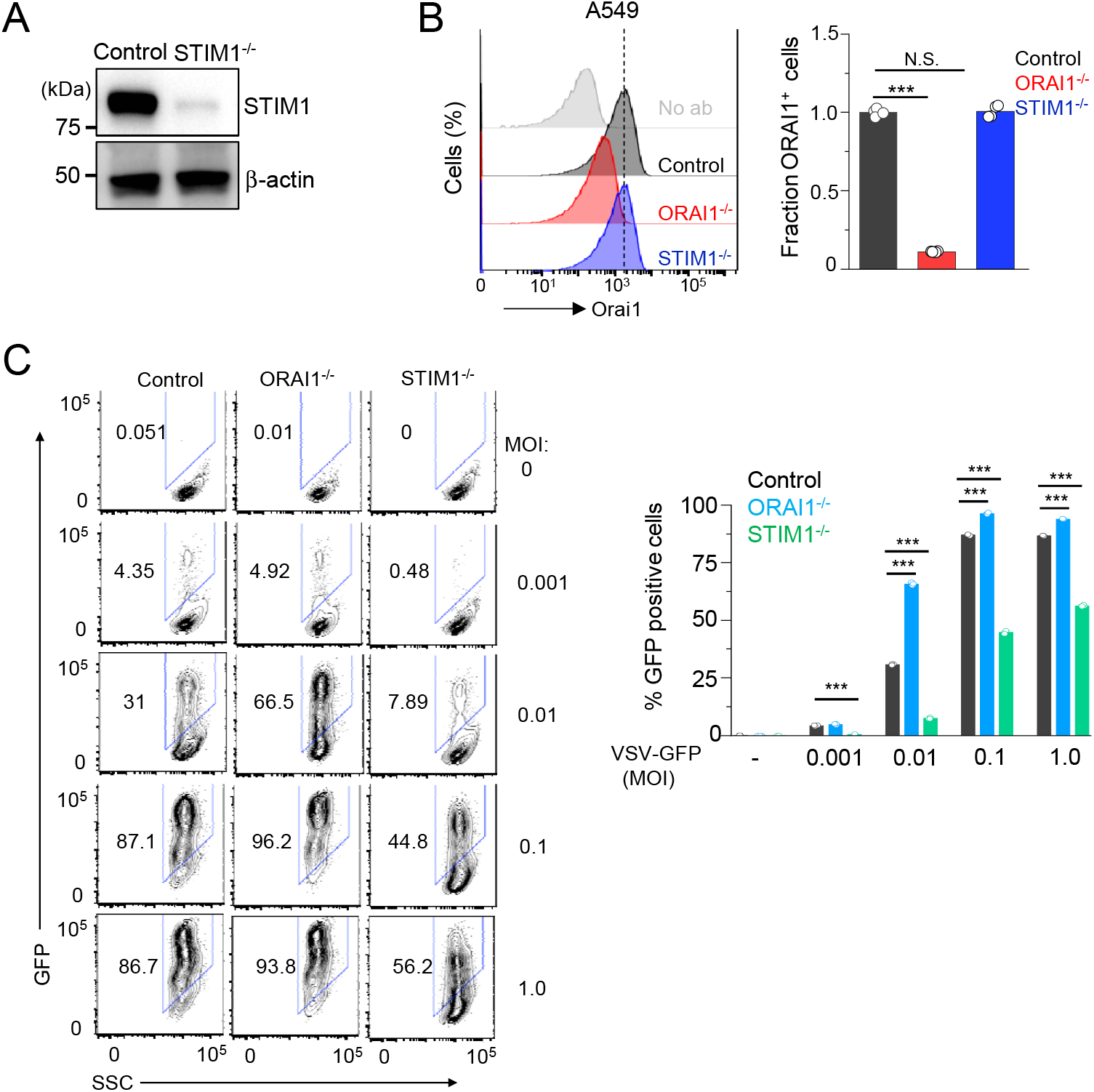
Loss of ORAI1 or STIM1 influences host resistance to vesicular stomatitis virus. (**A**) Representative immunoblot showing expression of STIM1 in lysates from control and *STIM1*^-/-^ A549 cells. β-actin – loading control. Data are representative of two independent experiments. (**B**) Representative histograms showing levels of total ORAI1 protein in control, *ORAI1*^-/-^, and *STIM1*^-/-^ A549 cells after permeabilization and intracellular staining with anti-ORAI1 antibody. The bar graph shows average (± s.e.m.) from three independent experiments. (**C**) Representative flow plots showing frequency of GFP-positive population in VSV-GFP-infected (at indicated MOIs for 20 hours) control, *ORAI1*^-/-^, or *STIM1*^-/-^ A549 cells. Bar graph (right) shows averaged frequency of VSV-GFP-positive populations from three independent experiments. \*\*\**P*<0.0005 (two-tailed *t* test).

## Discussion

The role of Ca^2+^ signaling mediated by ORAI1 and STIM1 in adaptive immune cells (e.g., T and B cells) is well established (Prakriya and Lewis, 2015; Srikanth and Gwack, 2013). However, the function of this pathway in the host-pathogen responses in innate immunity is poorly understood. A recent study had identified an important function of STIM1 in regulating the cytosolic DNA sensing pathway via direct interaction with STING (stimulator of interferon genes), independently from its role in Ca^2+^ signaling (Srikanth et al., 2019). The current study demonstrates a key role of ORAI1 in regulating tonic IFN-β levels, thereby host response to SARS-CoV-2 infection, by modulation of homeostatic cytoplasmic [Ca^2+^]. Collectively, these studies uncover novel functions of CRAC channel components in the innate immune system.

Loss of STIM1 provides strong resistance to infection with a DNA virus, herpes simplex virus type-1 (HSV-1), by enhancing IFN-I signaling (Srikanth et al., 2019). The observation of resistance to virus infections in *STIM1*^-/-^ cells has been extended to the current study using SARS-CoV-2. Abrogation of IFN-I signaling by co-deletion of IFNAR1 reduced baseline IFN-β secretion and rendered *STIM1*^-/-^ cells susceptible to SARS-CoV-2 infection, emphasizing the role of IFN-I in this resistance. Interestingly loss of ORAI1 did not affect host resistance to HSV-1, most likely because the STING signaling pathway, which is predominantly important for sensing DNA viruses, including HSV-1, was not impaired in *ORAI1*^-/-^ cells (Srikanth et al., 2019). However, loss of ORAI1 resulted in moderate susceptibility to VSV-GFP and very high susceptibility specifically to SARS-CoV-2 infection. SARS-CoV-2 shows high sensitivity to pre-treatment with type I or III IFNs, which profoundly reduces virus replication (Blanco-Melo et al., 2020; Lokugamage et al., 2020; Vanderheiden et al., 2020). Hence, reduction in tonic IFN-β levels may enhance the susceptibility of *ORAI1*^-/-^ cells especially to SARS-CoV-2 infection. It has been shown that in the absence of priming amounts of IFN-β, mouse embryonic fibroblasts do not produce other IFN-I, suggesting that IFN-β is a master regulator for all IFN-I activities and presumably those mediated by type III IFNs (Erlandsson et al., 1998; Takaoka et al., 2000). It has also been suggested that tonic IFN-β signaling is required to maintain adequate expression of STAT1 and 2, and accordingly, the absence of tonic IFN-β signaling reduces basal STAT expression (Gough et al., 2012), similar to our observation with *ORAI1*^-/-^ cells (**Fig. 4B**). Besides, loss of ORAI1 was shown to downregulate multiple pathways including, ISG antiviral signaling, TLR7/8 signaling as well as MyD88 signaling cascades involved in RNA sensing. Taken together, loss of tonic IFN-I signaling in combination with impaired function of other antiviral signaling cascades is likely to contribute towards the exquisite sensitivity of *ORAI1*^-/-^ cells to SARS-CoV-2.

The *IFNB* promoter contains four positive regulatory domains (PRDI-PRDIV), which are occupied by overlapping transcription factor complexes. Interferon regulatory factor 3 (IRF3) and IRF7 bind PRDI and III; the NF-κB RelA-p50 heterodimer binds PRDII; and the AP-1 heterodimer of ATF-2 and c-Jun binds PRDIV (Gough et al., 2012). The binding of each of these components in the correct orientation and location results in activation of the *IFNB* promoter in response to virus infection. Tonic IFN-β expression is independent of IRF3 and IRF7 but instead depends on the c-Jun and NF-κB p50 subunit. *ORAI1*^-/-^ cells showed reduced expression of multiple AP-1 family transcription factors, including ATF-2, FOS and JUN, as well as the NF-κB p50 subunit, which is likely to contribute to reduced baseline IFN-β levels in these cells. Notably, while SOCE was abolished in both *ORAI1*^-/-^ and *STIM1*^-/-^ cells, the baseline [Ca^2+^] was reduced only in *ORAI1*^-/-^ cells. Based on the previous finding that STIM2, that has a lower binding affinity to Ca^2+^ than STIM1, is crucial for regulation of homeostatic Ca^2+^ levels (Brandman et al., 2007), it is expected that *STIM2*^-/-^ cells may have lower tonic IFN-I signaling, thereby high susceptibility to SARS-CoV-2 infection similar to *ORAI1*^-/-^ cells. Another notable difference between ORAI1 and STIM1 deficiency is differential expression of IFN-β and IL-6. *STIM1*^-/-^ cells showed enhanced expression of both these cytokines, in consistence with the observation that activation of the STING pathway upregulates expression of both IFN-β and IL-6 (Decout et al., 2021; Srikanth et al., 2019). However, *ORAI1*^-/-^ cells had normal expression of IL-6, but still showed high susceptibility to SARS-CoV-2 infection, suggesting that the protective effect of STIM1 deficiency is predominantly derived from increased levels of IFN-β. Therefore, these results obtained from the analysis of *ORAI1*^-/-^ and *STIM1*^-/-^ cells cohesively suggest that the baseline IFN-I signaling is a crucial determinant for the degree of host resistance to SARS-CoV-2.

In summary, we examined the role of CRAC channel components, ORAI1 and STIM1, in host resistance to SARS-CoV-2 infection. *STIM1*^-/-^ cells showed remarkable resistance to SARS-CoV-2 infection supporting previous studies of the role of elevated baseline IFN-I levels (Srikanth et al., 2019). Loss of ORAI1 severely impaired the tonic IFN-I levels, thereby reducing resistance to SARS-CoV-2 infection. Mechanistically, ORAI1-mediated Ca^2+^ signaling affects expression of various transcription factors including *FOS*, *JUN*, *ATF2* and *MEF2C* that are involved in regulating multiple host defense pathways. The key finding of this study is that the baseline IFN-β level regulated by ORAI1-mediated Ca^2+^ signaling is critical for priming the cellular antiviral state, determining host resistance to SARS-CoV-2 infection. A recent phase 2 clinical trial of inhaled, nebulized IFN-β to treat people with COVID-19 resulted in accelerated recovery, supporting localized delivery of IFN-I as a potential prophylactic and therapeutic agent against COVID-19 (Monk et al., 2021). It is likely that the therapeutic effect of IFN-I targeting SARS-CoV-2 infection may be partly derived from boosting tonic IFN-I signaling. The current study suggests that increasing baseline cytoplasmic [Ca^2+^] by enhancing ORAI1 activity is likely to boost cellular antiviral defense mechanisms to multiple RNA viruses including SARS-CoV-2.

## Materials and Methods

### Chemicals and Antibodies

Fura 2-AM (F1221) was purchased from Thermofisher Scientific. Thapsigargin and cyclosporin A were purchased from EMD Millipore. SARS-CoV Spike (S) antibodies - polyclonal anti-SARS coronavirus (BEI Resources: NR-10361 antiserum, Guinea Pig) was used for immunoblotting and monoclonal anti-SARS-CoV S protein antibody (BEI Resources: NR-616, similar to 240C) was used for immunocytochemistry. Antibodies for detection of human STIM1 (5668S) was purchased from Cell Signaling Technologies, that for detection of interferon alpha and beta receptor 1 (IFNAR-1) was purchased from Leinco Technologies, Inc. (I-400), that for detection of surface ORAI1 (ACC-060) was purchased from Alomone labs and that for detection of β-actin (sc-47778), was obtained from Santa Cruz Biotechnology.

### Plasmids, sgRNAs and cells

Human ACE2 coding sequence was cloned into a lentiviral vector as described (Garcia et al., 2021). sgRNAs targeting human *ORAI1* (Forward primer: *CACCGGATCGGCCAGAGTTACTCC*; Reverse primer: *AAACCGGAGTAACTCTGGCCGATCC*) and human *STIM1* (Forward primer: *CACCGTGAGGATAAGCTCATCAGCG*; Reverse Primer: *AAACCGCTGATGAGCTTATCCTCAC*) genes were subcloned into pLentiguide puro (Addgene). sgRNAs targeting human interferon alpha and beta receptor subunit 1 (IFNAR1) subcloned into pLentiCRISPR v2 was purchased from GenScript (catalog # IFNAR1 crRNA 1; gRNA target sequence *GGCGTGTTTCCAGACTGTTT*). Vero E6, HEK293 and A549 cells were obtained from the American Type Culture Collection center (ATCC, Manassas, VA). Vero cells were cultured in EMEM growth media (ATCC) supplemented with 10% (v/v) fetal bovine serum (FBS, Hyclone) and Penicillin/Streptomycin (Mediatech) at 37°C and 5% CO_2_. HEK293 and A549 cells were cultured in complete DMEM (Mediatech) supplemented with 10% (v/v) FBS (Hyclone), 2 mM L-glutamine (Mediatech), 10 mM HEPES (Mediatech) and Penicillin/Streptomycin (Mediatech) at 37°C and 5% CO_2_.

### Virus

SARS-CoV-2 (USA-WA1/2020) was obtained from BEI Resources of National Institute of Allergy and Infectious Diseases (NIAID). All the studies involving live virus was conducted in UCLA BSL3 high-containment facility. SARS-CoV-2 was passaged once in Vero E6 cells and viral stocks were aliquoted and stored at −80°C. Virus titer was measured in Vero E6 cells by established plaque assay.

### Lentiviral transduction of HEK293 and A549 cells

To generate HEK293-ACE2 cells, HEK293T cells were transfected with plasmid encoding ACE2 and packaging vectors (pMD2.G and psPAX2, Addgene) using calcium phosphate transfection method. Culture supernatants were harvested at 48 and 72 hours post transfection, filtered using a 0.45 μ PVDF filter and used for infection of HEK293 cells together with polybrene (8 μg/ml) using the spin-infection method. Transduced cells were selected with puromycin (1 μg/ml) 48 hours post second infection. For generation of *ORAI1*^-/-^ and *STIM1*^-/-^ cells, HEK293T cells were transfected with plasmids encoding sgRNA and packaging vectors as described above. Lentiviruses encoding Cas9 were generated using the same technique. Culture supernatants harvested at 48 and 72 hours post transfection were used for infection (50% of Cas9-encoding virus + 50% of sgRNA-encoding virus) of HEK293-ACE2 or A549 cells together with polybrene (8 μg/ml) using the spin-infection method. Cells were selected with puromycin (1 μg/ml) and blasticidin (5 μg/ml) 48 hours post second infection.

### Endogenous ORAI1 staining and analysis

For total endogenous ORAI1 detection, control, STIM1 KO and ORAI1 KO HEK293 or A549 cells (0.5 × 10^6^) were fixed with 4% PFA at room temperature for 15 mins, followed by permeabilization with PBS + 0.5% Igepal CA-630 and incubated with 1 μg unlabeled anti-ORAI1 antibody (ACC-060, Alomone labs) in PBS + 2% FBS + 0.5% Igepal CA-630. Negative control sample did not contain the primary Ab. Cells were subsequently treated with FITC-conjugated secondary antibody, washed and data was acquired on a BD Fortessa flow cytometer (BD Biosciences) and analyzed using FlowJo software (Treestar Inc).

### Single-cell Ca^2+^ imaging and immunofluorescence analysis

HEK293-ACE and A549 cells were grown overnight on coverslips and loaded with 1 μM Fura 2-AM for 40 min at 25°C for imaging. Intracellular [Ca^2+^]_i_ measurements were performed using essentially the same methods as previously described (Srikanth et al., 2010b). Briefly, microscopy was performed using an Olympus IX2 illumination system mounted on an Olympus IX51 inverted microscope using previously described methods (Srikanth et al., 2010a). Acquisition and image analysis were performed using Slidebook (Intelligent Imaging Innovations, Inc.) software and graphs were plotted using Origin2020 (OriginLab). For immunofluorescence analysis, uninfected or SARS-CoV-2-infected cells were fixed for 20 mins with ice-cold methanol, permeabilized in buffer containing PBS + 0.2% Triton X-100, blocked with same buffer containing 2.5% goat serum, 2.5% donkey serum and 2% BSA. Cells were stained overnight in the blocking buffer with primary antibodies washed and treated with secondary antibody for 1 h, stained for DAPI in permeabilization solution and visualized using 40X oil immersion lens and imaged using Slidebook 6.0 software (Intelligent Imaging Innovations, Inc.). Images were processed for enhancement of brightness or contrast using Slidebook 6.0 software.

### SARS-CoV-2 Infection

Control, *ORAI1*^-/-^, or *STIM1*^-/-^ HEK293-ACE2 cells were seeded at 1 x 10^5^ cells per well in 0.4 ml volumes using a 48-well plate, or 5 x 10^4^ cells per well in 0.2 ml volume in a 96-well plate. The following day, viral inoculum (MOI of 0.1 or 1.0 as indicated) was prepared using serum free media. The spent media from each well was removed and 100 μl of prepared inoculum was added onto cells. For mock infection, serum free media (100 μl/well) alone was added. The inoculated plates were incubated for 1 hour at 37 °C with 5% CO_2_. The inoculum was spread by gently tilting the plate sideways every 15 minutes. At the end of incubation, the inoculum was replaced with serum supplemented media. At 20 hours post infection (hpi) the cells were either fixed with methanol for immunocytochemistry (ICC) analysis or harvested in TriZol reagent (ThermoFisher) for RNA-seq and RT-PCR analysis or lysed in RIPA lysis buffer for immunoblotting as described below.

### Plaque Assay

Viral titers were determined by plaque assay on Vero E6 cells. Vero E6 cells were seeded in 12-well plates for 24 hours. Virus-containing supernatants were 10-fold serially diluted in PBS. The growth medium was removed from the cells, cells were washed once with PBS, and diluted supernatants were added (150 μl/well). After 30 min inoculation, an overlay medium (double-concentrated minimal essential medium (MEM; supplemented with 2 % L-glumine, 2 % Pen/Strep, 0.4 % bovine serum albumin (BSA)), mixed 1:1 with 2.5 % avicel solution (prepared in ddH2O)) was added to the cells (1.5 ml/well). Then, cells were incubated for 72 hours at 37 °C. After 72 hours, the overlay medium was removed from the cells, and following a washing step with 1x PBS the cells were fixed with 4 % paraformaldehyde (PFA) for 30 min at 4 °C. Subsequently, the cells were counterstained with crystal violet solution to visualize the virus-induced plaques in the cell layer. The number of plaques at a given dilution were counted to calculate the viral titers as plaque-forming units (PFU/ml).

### RNA sample preparation and RT-qPCR

Total RNA from cells harvested in TRIzol Reagent (ThermoFisher) was isolated using the Direct-zol RNA isolation kit (Zymo Research). RNA quantity and quality were confirmed with a NanoDrop ND-1000 spectrophotometer. cDNA was synthesized using 0.5-1.0 μg of total RNA using random hexamers and oligo(dT) primers and qScript cDNA Supermix (Quantabio). Real-time quantitative PCR was performed using iTaq Universal SYBR Green Supermix (Bio-Rad) and an iCycler IQ5 system (Bio-Rad) using gene-specific primers. Briefly, amplification was performed using 20 μl volume reactions in a 96-well plate format with the following conditions: 95°C for 30 sec for polymerase activation and cDNA denaturation, then 40 cycles at 95°C for 15 seconds and 60°C for 1 minute, with a melt-curve analysis at 65-95°C and 0.5°C increments at 2-5 seconds/step. Threshold cycles (C_T_) for all the candidate genes were normalized to those of 36B4 to obtain ΔC_T_. The specificity of primers was examined by melt-curve analysis and agarose gel electrophoresis of PCR products. Primer sequences are: *SARS-CoV-2*: (Forward: *GACCCCAAAATCAGCGAAAT*, Reverse: *TCTGGTTACTGCCAGTTGAATCTG*), human *ATF2* (Forward: *GCACAGCCCACATCAGCTATT*, Reverse: *GGTGCCTGGGTGATTACAGT*); human *FOS* (Forward: *GGGGCAAGGTGGAACAGTTAT*, Reverse: *CCGCTTGGAGTGTATCAGTCA*); human *MEF2C* (Forward: *GTATGGCAATCCCCGAAACT*, Reverse: *ATCGTATTCTTGCTGCCTGG*), and human *36B4* (Forward: *AGATGCAGCAGATCCGCAT*, Reverse: *GTTCTTGCCCATCAGCACC*).

### Western Blot analysis

Cells were lysed in RIPA lysis buffer (50 mM Tris pH 7.4, 1% NP-40, 0.25% sodium deoxycholate, 1 mM EDTA, 150 mM NaCl, 1 mM Na3VO4, 20 mM NaF, 1mM PMSF, 2 mg ml^-1^ aprotinin, 2 mg ml^-1^ leupeptin and 0.7 mg ml^-1^ pepstatin) for 1 h with intermittent vortexing and centrifuged to remove debris. Samples were separated on 10% SDS-PAGE. Proteins were transferred to nitrocellulose membranes and subsequently analyzed by immunoblotting with relevant antibodies. Chemiluminescence images were acquired using an Image reader LAS-3000 LCD camera (FujiFilm).

### Cytokine measurement by ELISA

ELISA was performed on cell culture supernatants from indicated control or SARS-CoV-2-infected cells for detection of IFN-β (PBL Assay Science, #41410) or IL-6 (ThermoFisher Scientific, # 88-7066-22) using manufacturer’s instructions.

### RNA sequencing and data analysis

RNA was extracted with Direct-zol RNA Mini prep kit (Zymo Research). Libraries for RNA-Seq were prepared with TruSeq Stranded mRNA Library Prep Kit (Illumina). The workflow consisted of mRNA enrichment and fragmentation. Cleaved RNA fragments were copied into first strand cDNA using reverse transcriptase and random primers. Strand specificity is achieved by replacing dTTP with dUTP, followed by second strand cDNA synthesis using DNA Polymerase I and RNase H. cDNA generation is followed by A-tailing, adaptor ligation and PCR amplification. Different adaptors were used for multiplexing samples in one lane. Sequencing was performed on Illumina HiSeq 3000 for SE 1×50 run. Data quality check was done on Illumina SAV. Demultiplexing was performed with Illumina Bcl2fastq v2.19.1.403 software. Raw reads for Orai1KO and STIM1KO SARS-CoV-2 infected samples (Orai1KO-SARS-CoV-2-3_S18 and STIM1KO-SARS-CoV-2-1_S19) were sampled down to 45 million reads before mapping. Illumina reads from all the samples were mapped to human and SARS-CoV-2 reference genomes by STAR v2.27a (Dobin et al., 2013) and read counts per gene were quantified using human Ensembl GRCh38.98 and SARS-CoV-2 GTF file to generate raw reads counts. Differential expression analysis was performed using DESeq2 v1.28.1 in R v4.0.3 (Love et al., 2014). Median of ratios method was used to normalize expression counts for each gene in all the experimental samples. Each gene in the samples was fitted into a negative binomial generalized linear model. Differentially expressed gene (DEG) candidates were considered if they were supported by a false discovery rate (FDR) P < 0.01. Unsupervised principal component analysis was performed using pcaExplorer (Marini and Binder, 2019) in R v4.0.3 (Love et al., 2014). Reactome pathway analysis was performed for DEGs using human all genes as reference data set in the Reactome v65 (Jassal et al., 2020) implemented in PANTHER. Reactome pathways were only considered if they were supported by FDR P < 0.05. The ggplot2 v3.3.2 in R and Prism GraphPad v8.4.3 were used to generate figures. The shinyheatmap web interface was used to prepare heat maps (Khomtchouk et al., 2017). The genes in the DEGs directly regulated by transcription factors (FOS, ATF2, MEF2C) were identified based on binding profiles of all public ChIP-seq data for particular gene loci from the ChIP-Atlas database (Oki et al., 2018). The regulated or target genes were accepted if the peak-call intervals of a given gene overlapped with a transcription start site (TSS) +/-1 Kb. RNA sequencing data are deposited in NCBI GEO under the accession number GSE173707.

### Statistical analysis

Statistical analysis was carried out using two-tailed Student’s t-test. Differences were considered significant when *p* values were <0.05. The hypergeometric test in R v4.0.3 was used to measure the statistical significance of the enriched genes in the DEGs regulated by transcription factors (FOS, ATF2, MEF2C) compared to regulated genes in the background gene sets of the human reference genome.

## ACKNOWLEDGEMENTS

The authors thank Barbara Dillon, UCLA High-Containment Program Director for BSL3 work. The authors would also like to thank Spyridon Hasiakos (UCLA Oral Biology Graduate Student) and Shawn Cokus (UCLA Collaboratory Fellow) for suggestions about statistical analysis of transcription factor enrichment data. Flow cytometry was performed in the UCLA Jonsson Comprehensive Cancer Center (JCCC) and Center for AIDS Research Flow Cytometry Core Facility that is supported by National Institutes of Health awards P30 CA016042 and 5P30 AI028697, and by the JCCC, the UCLA AIDS Institute, the David Geffen School of Medicine at UCLA, the UCLA Chancellor’s Office, and the UCLA Vice Chancellor’s Office of Research. This work was supported by the National Institute of Health grants AI146352 (S.S.), EY032149 (V.A.), AI083432, AI146615, AI147063, AI149236 (Y.G.), Broad Stem cell Research Center institutional awards OCRC 20-76 (S.S.) and OCRC 20-15 (V.A.), and the California Institute for Regenerative Medicine Tran Award, TRAN1COVID19-11975 (V.A.). A.R. is supported by the Tata Institute for Genetics and Society.

## AUTHOR CONTRIBUTIONS

S.S. and Y.G. conceptualized the study. S.S. generated all the cells and performed Ca^2+^ imaging experiments. B.B.W. performed all the molecular biology and flow cytometry experiments and A.R. performed the RNA-seq data and bioinformatics analysis. G.G. and V.A. generated the SARS-CoV-2 virus and performed all infection experiments. S.S. wrote the manuscript with input from all authors.

## References

Ban, N., Yamada, Y., Someya, Y., Ihara, Y., Adachi, T., Kubota, A., Watanabe, R., Kuroe, A., Inada, A., Miyawaki, K., et al. (2000). Activating transcription factor-2 is a positive regulator in CaM kinase IV-induced human insulin gene expression. Diabetes 49, 1142–1148.

Bastard, P., Rosen, L.B., Zhang, Q., Michailidis, E., Hoffmann, H.H., Zhang, Y., Dorgham, K., Philippot, Q., Rosain, J., Beziat, V., et al. (2020). Autoantibodies against type I IFNs in patients with life-threatening COVID-19. Science 370.

Blanco-Melo, D., Nilsson-Payant, B.E., Liu, W.C., Uhl, S., Hoagland, D., Moller, R., Jordan, T.X., Oishi, K., Panis, M., Sachs, D., et al. (2020). Imbalanced Host Response to SARS-CoV-2 Drives Development of COVID-19. Cell 181, 1036–1045 e1039.

Brandman, O., Liou, J., Park, W.S., and Meyer, T. (2007). STIM2 is a feedback regulator that stabilizes basal cytosolic and endoplasmic reticulum Ca2+ levels. Cell 131, 1327–1339.

Chen, G., Wu, D., Guo, W., Cao, Y., Huang, D., Wang, H., Wang, T., Zhang, X., Chen, H., Yu, H., et al. (2020). Clinical and immunological features of severe and moderate coronavirus disease 2019. J Clin Invest 130, 2620–2629.

Crotta, S., Davidson, S., Mahlakoiv, T., Desmet, C.J., Buckwalter, M.R., Albert, M.L., Staeheli, P., and Wack, A. (2013). Type I and type III interferons drive redundant amplification loops to induce a transcriptional signature in influenza-infected airway epithelia. PLoS Pathog 9, e1003773.

Daniloski, Z., Jordan, T.X., Wessels, H.H., Hoagland, D.A., Kasela, S., Legut, M., Maniatis, S., Mimitou, E.P., Lu, L., Geller, E., et al. (2021). Identification of Required Host Factors for SARS-CoV-2 Infection in Human Cells. Cell 184, 92–105 e116.

Decout, A., Katz, J.D., Venkatraman, S., and Ablasser, A. (2021). The cGAS-STING pathway as a therapeutic target in inflammatory diseases. Nat Rev Immunol.

Dobin, A., Davis, C.A., Schlesinger, F., Drenkow, J., Zaleski, C., Jha, S., Batut, P., Chaisson, M., and Gingeras, T.R. (2013). STAR: ultrafast universal RNA-seq aligner. Bioinformatics 29, 15–21.

Erlandsson, L., Blumenthal, R., Eloranta, M.L., Engel, H., Alm, G., Weiss, S., and Leanderson, T. (1998). Interferon-beta is required for interferon-alpha production in mouse fibroblasts. Curr Biol 8, 223–226.

Garcia, G., Jr., Sharma, A., Ramaiah, A., Sen, C., Purkayastha, A., Kohn, D.B., Parcells, M.S., Beck, S., Kim, H., Bakowski, M.A., et al. (2021). Antiviral drug screen identifies DNA-damage response inhibitor as potent blocker of SARS-CoV-2 replication. Cell Rep, 108940.

Gough, D.J., Messina, N.L., Clarke, C.J., Johnstone, R.W., and Levy, D.E. (2012). Constitutive type I interferon modulates homeostatic balance through tonic signaling. Immunity 36, 166–174.

Hoffmann, H.H., Schneider, W.M., Rozen-Gagnon, K., Miles, L.A., Schuster, F., Razooky, B., Jacobson, E., Wu, X., Yi, S., Rudin, C.M., et al. (2021). TMEM41B Is a Pan-flavivirus Host Factor. Cell 184, 133–148 e120.

Iwasaki, A., and Medzhitov, R. (2015). Control of adaptive immunity by the innate immune system. Nat Immunol 16, 343–353.

Jassal, B., Matthews, L., Viteri, G., Gong, C., Lorente, P., Fabregat, A., Sidiropoulos, K., Cook, J., Gillespie, M., Haw, R., et al. (2020). The reactome pathway knowledgebase. Nucleic Acids Res 48, D498–D503.

Khomtchouk, B.B., Hennessy, J.R., and Wahlestedt, C. (2017). shinyheatmap: Ultra fast low memory heatmap web interface for big data genomics. PLoS One 12, e0176334.

Lazear, H.M., Schoggins, J.W., and Diamond, M.S. (2019). Shared and Distinct Functions of Type I and Type III Interferons. Immunity 50, 907–923.

Lesch, A., Hui, X., Lipp, P., and Thiel, G. (2015). Transient receptor potential melastatin-3 (TRPM3)-induced activation of AP-1 requires Ca2+ ions and the transcription factors c-Jun, ATF2, and ternary complex factor. Mol Pharmacol 87, 617–628.

Li, W., Moore, M.J., Vasilieva, N., Sui, J., Wong, S.K., Berne, M.A., Somasundaran, M., Sullivan, J.L., Luzuriaga, K., Greenough, T.C., et al. (2003). Angiotensin-converting enzyme 2 is a functional receptor for the SARS coronavirus. Nature 426, 450–454.

Lokugamage, K.G., Hage, A., de Vries, M., Valero-Jimenez, A.M., Schindewolf, C., Dittmann, M., Rajsbaum, R., and Menachery, V.D. (2020). Type I Interferon Susceptibility Distinguishes SARS-CoV-2 from SARS-CoV. J Virol 94.

Love, M.I., Huber, W., and Anders, S. (2014). Moderated estimation of fold change and dispersion for RNA-seq data with DESeq2. Genome Biol 15, 550.

Marini, F., and Binder, H. (2019). pcaExplorer: an R/Bioconductor package for interacting with RNA-seq principal components. BMC Bioinformatics 20, 331.

McKinsey, T.A., Zhang, C.L., and Olson, E.N. (2002). MEF2: a calcium-dependent regulator of cell division, differentiation and death. Trends Biochem Sci 27, 40–47.

Monk, P.D., Marsden, R.J., Tear, V.J., Brookes, J., Batten, T.N., Mankowski, M., Gabbay, F.J., Davies, D.E., Holgate, S.T., Ho, L.P., et al. (2021). Safety and efficacy of inhaled nebulised interferon beta-1a (SNG001) for treatment of SARS-CoV-2 infection: a randomised, double-blind, placebo-controlled, phase 2 trial. Lancet Respir Med 9, 196–206.

Ng, S.W., Nelson, C., and Parekh, A.B. (2009). Coupling of Ca(2+) microdomains to spatially and temporally distinct cellular responses by the tyrosine kinase Syk. J Biol Chem 284, 24767–24772.

Nomura, A., Yokoe, S., Tomoda, K., Nakagawa, T., Martin-Romero, F.J., and Asahi, M. (2020). Fluctuation in O-GlcNAcylation inactivates STIM1 to reduce store-operated calcium ion entry via down-regulation of Ser(621) phosphorylation. J Biol Chem 295, 17071–17082.

Oki, S., Ohta, T., Shioi, G., Hatanaka, H., Ogasawara, O., Okuda, Y., Kawaji, H., Nakaki, R., Sese, J., and Meno, C. (2018). ChIP-Atlas: a data-mining suite powered by full integration of public ChIP-seq data. EMBO Rep 19.

Prakriya, M., and Lewis, R.S. (2015). Store-Operated Calcium Channels. Physiol Rev 95, 1383–1436.

Schneider, W.M., Luna, J.M., Hoffmann, H.H., Sanchez-Rivera, F.J., Leal, A.A., Ashbrook, A.W., Le Pen, J., Ricardo-Lax, I., Michailidis, E., Peace, A., et al. (2021). Genome-Scale Identification of SARS-CoV-2 and Pancoronavirus Host Factor Networks. Cell 184, 120–132 e114.

Schulz, K.S., and Mossman, K.L. (2016). Viral Evasion Strategies in Type I IFN Signaling - A Summary of Recent Developments. Front Immunol 7, 498.

Srikanth, S., and Gwack, Y. (2013). Orai1-NFAT signalling pathway triggered by T cell receptor stimulation. Mol Cells 35, 182–194.

Srikanth, S., Jung, H.J., Kim, K.D., Souda, P., Whitelegge, J., and Gwack, Y. (2010a). A novel EF-hand protein, CRACR2A, is a cytosolic Ca2+ sensor that stabilizes CRAC channels in T cells. Nat Cell Biol 12, 436–446.

Srikanth, S., Jung, H.J., Ribalet, B., and Gwack, Y. (2010b). The intracellular loop of Orai1 plays a central role in fast inactivation of Ca2+ release-activated Ca2+ channels. J Biol Chem 285, 5066–5075.

Srikanth, S., Woo, J.S., Wu, B., El-Sherbiny, Y.M., Leung, J., Chupradit, K., Rice, L., Seo, G.J., Calmettes, G., Ramakrishna, C., et al. (2019). The Ca(2+) sensor STIM1 regulates the type I interferon response by retaining the signaling adaptor STING at the endoplasmic reticulum. Nat Immunol 20, 152–162.

Takaoka, A., Mitani, Y., Suemori, H., Sato, M., Yokochi, T., Noguchi, S., Tanaka, N., and Taniguchi, T. (2000). Cross talk between interferon-gamma and -alpha/beta signaling components in caveolar membrane domains. Science 288, 2357–2360.

tenOever, B.R. (2016). The Evolution of Antiviral Defense Systems. Cell Host Microbe 19, 142–149.

Vanderheiden, A., Ralfs, P., Chirkova, T., Upadhyay, A.A., Zimmerman, M.G., Bedoya, S., Aoued, H., Tharp, G.M., Pellegrini, K.L., Manfredi, C., et al. (2020). Type I and Type III Interferons Restrict SARS-CoV-2 Infection of Human Airway Epithelial Cultures. J Virol 94.

Wang, R., Simoneau, C.R., Kulsuptrakul, J., Bouhaddou, M., Travisano, K.A., Hayashi, J.M., Carlson-Stevermer, J., Zengel, J.R., Richards, C.M., Fozouni, P., et al. (2021). Genetic Screens Identify Host Factors for SARS-CoV-2 and Common Cold Coronaviruses. Cell 184, 106–119 e114.

Wei, J., Alfajaro, M.M., DeWeirdt, P.C., Hanna, R.E., Lu-Culligan, W.J., Cai, W.L., Strine, M.S., Zhang, S.M., Graziano, V.R., Schmitz, C.O., et al. (2021). Genome-wide CRISPR Screens Reveal Host Factors Critical for SARS-CoV-2 Infection. Cell 184, 76–91 e13.

Xia, H., and Shi, P.Y. (2020). Antagonism of Type I Interferon by Severe Acute Respiratory Syndrome Coronavirus 2. J Interferon Cytokine Res 40, 543–548.

Xie, M., and Chen, Q. (2020). Insight into 2019 novel coronavirus - An updated interim review and lessons from SARS-CoV and MERS-CoV. Int J Infect Dis 94, 119–124.

Yoast, R.E., Emrich, S.M., Zhang, X., Xin, P., Johnson, M.T., Fike, A.J., Walter, V., Hempel, N., Yule, D.I., Sneyd, J., et al. (2020). The native ORAI channel trio underlies the diversity of Ca(2+) signaling events. Nat Commun 11, 2444.

Zhang, Q., Bastard, P., Liu, Z., Le Pen, J., Moncada-Velez, M., Chen, J., Ogishi, M., Sabli, I.K.D., Hodeib, S., Korol, C., et al. (2020). Inborn errors of type I IFN immunity in patients with life-threatening COVID-19. Science 370.

