## Supplemental Figure Legends for "ORAI1 establishes resistance to SARS-CoV-2 infection by regulating tonic type I interferon signaling"

**Supplementary Figure 1. Similar expression of ACE2 in control, *ORAI1*$^{-/-}$, and *STIM1*$^{-/-}$ HEK293-ACE2 cells.**

(**A**) Representative epifluorescence images of indicated HEK293-ACE2 cells stained to check expression of ACE2. DAPI – nuclear stain. Data are representative of two independent experiments.

(**B**) Representative immunoblot showing expression of ACE2 in lysates from control, *ORAI1*$^{-/-}$, or *STIM1*$^{-/-}$ HEK293-ACE2 cells. β-actin – loading control. Data are representative of two independent experiments.

**Supplementary Figure 2. Principal component analysis of RNA-seq data.**

Principal component analysis of RNA-seq comparing data obtained from HEK293-ACE2 control samples with *ORAI1*$^{-/-}$ or *STIM1*$^{-/-}$ cells as indicated under mock conditions or 20 hours after infection with SARS-CoV-2. Each dot represents a biological replicate.

**Supplementary Figure 3. Heat map illustrating z-scores for host factors essential for SARS-CoV-2 replication, differentially expressed between control and *ORAI1*$^{-/-}$ HEK293-ACE2 cells.**

The gene list was derived from indicated references (Daniloski et al., 2021; Hoffmann et al., 2021; Schneider et al., 2021; Wang et al., 2021; Wei et al., 2021).

### References:

Daniloski, Z., Jordan, T.X., Wessels, H.H., Hoagland, D.A., Kasela, S., Legut, M., Maniatis, S., Mimitou, E.P., Lu, L., Geller, E., *et al.* (2021). Identification of Required Host Factors for SARS-CoV-2 Infection in Human Cells. Cell *184*, 92-105 e116.

Hoffmann, H.H., Schneider, W.M., Rozen-Gagnon, K., Miles, L.A., Schuster, F., Razooky, B., Jacobson, E., Wu, X., Yi, S., Rudin, C.M., *et al.* (2021). TMEM41B Is a Pan-flavivirus Host Factor. Cell *184*, 133-148 e120.

Schneider, W.M., Luna, J.M., Hoffmann, H.H., Sanchez-Rivera, F.J., Leal, A.A., Ashbrook, A.W., Le Pen, J., Ricardo-Lax, I., Michailidis, E., Peace, A., *et al.* (2021). Genome-Scale Identification of SARS-CoV-2 and Pan-coronavirus Host Factor Networks. Cell *184*, 120-132 e114.

Wang, R., Simoneau, C.R., Kulsuptrakul, J., Bouhaddou, M., Travisano, K.A., Hayashi, J.M., Carlson-Stevermer, J., Zengel, J.R., Richards, C.M., Fozouni, P., *et al.* (2021). Genetic Screens Identify Host Factors for SARS-CoV-2 and Common Cold Coronaviruses. Cell *184*, 106-119 e114.

Wei, J., Alfajaro, M.M., DeWeirdt, P.C., Hanna, R.E., Lu-Culligan, W.J., Cai, W.L., Strine, M.S., Zhang, S.M., Graziano, V.R., Schmitz, C.O., *et al.* (2021). Genome-wide CRISPR Screens Reveal Host Factors Critical for SARS-CoV-2 Infection. Cell *184*, 76-91 e13.
