## Supplemental Figures 1-3 for "ORAI1 establishes resistance to SARS-CoV-2 infection by regulating tonic type I interferon signaling"

### Slide 1
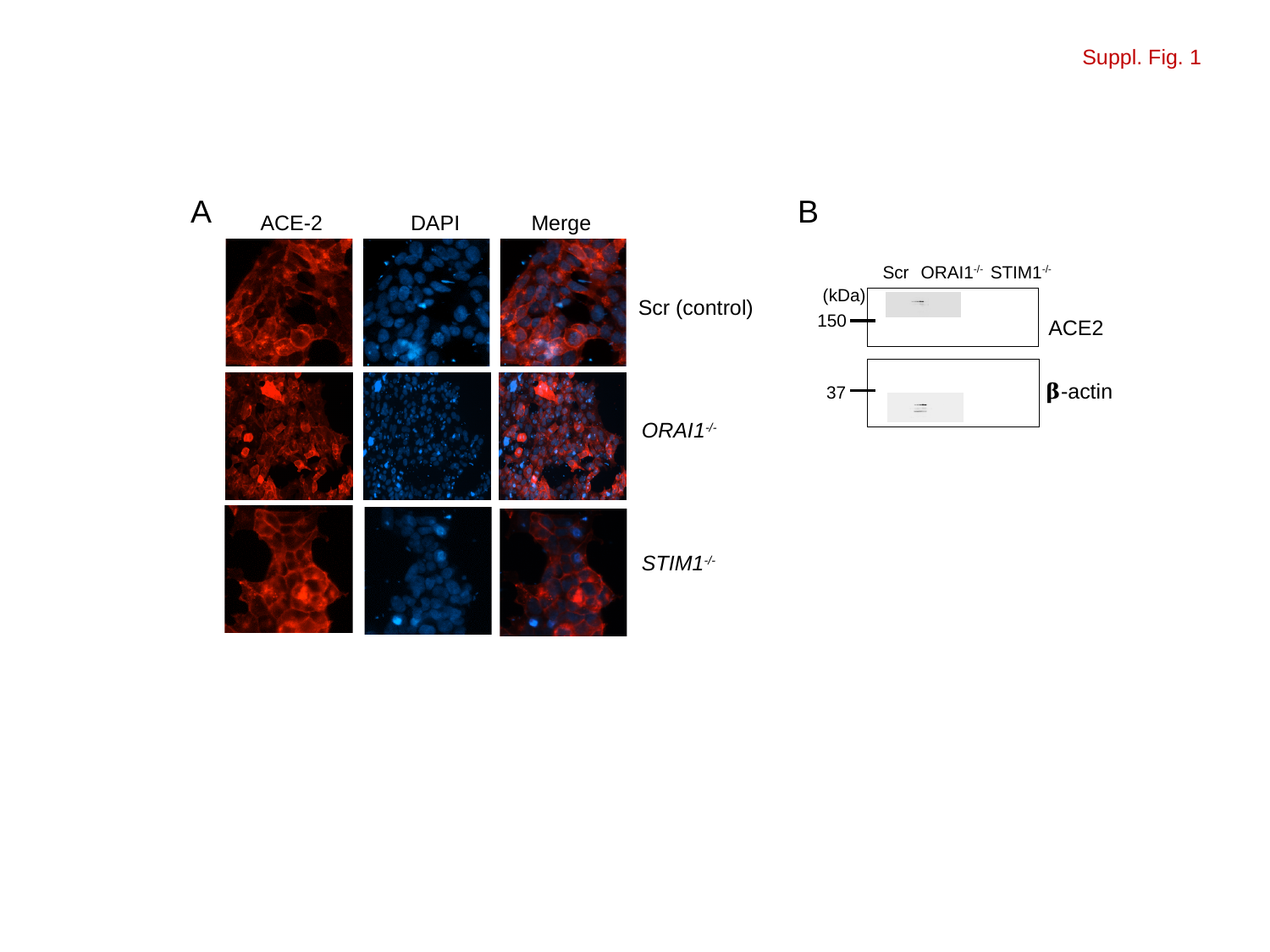

Suppl. Fig. 1
A
B
ACE-2
DAPI
Merge
Scr
ORAI1^-/-^
STIM1^-/-^
(kDa)
Scr (control)
150
ACE2
𝛃-actin
37
ORAI1^-/-^
STIM1^-/-^

### Slide 2
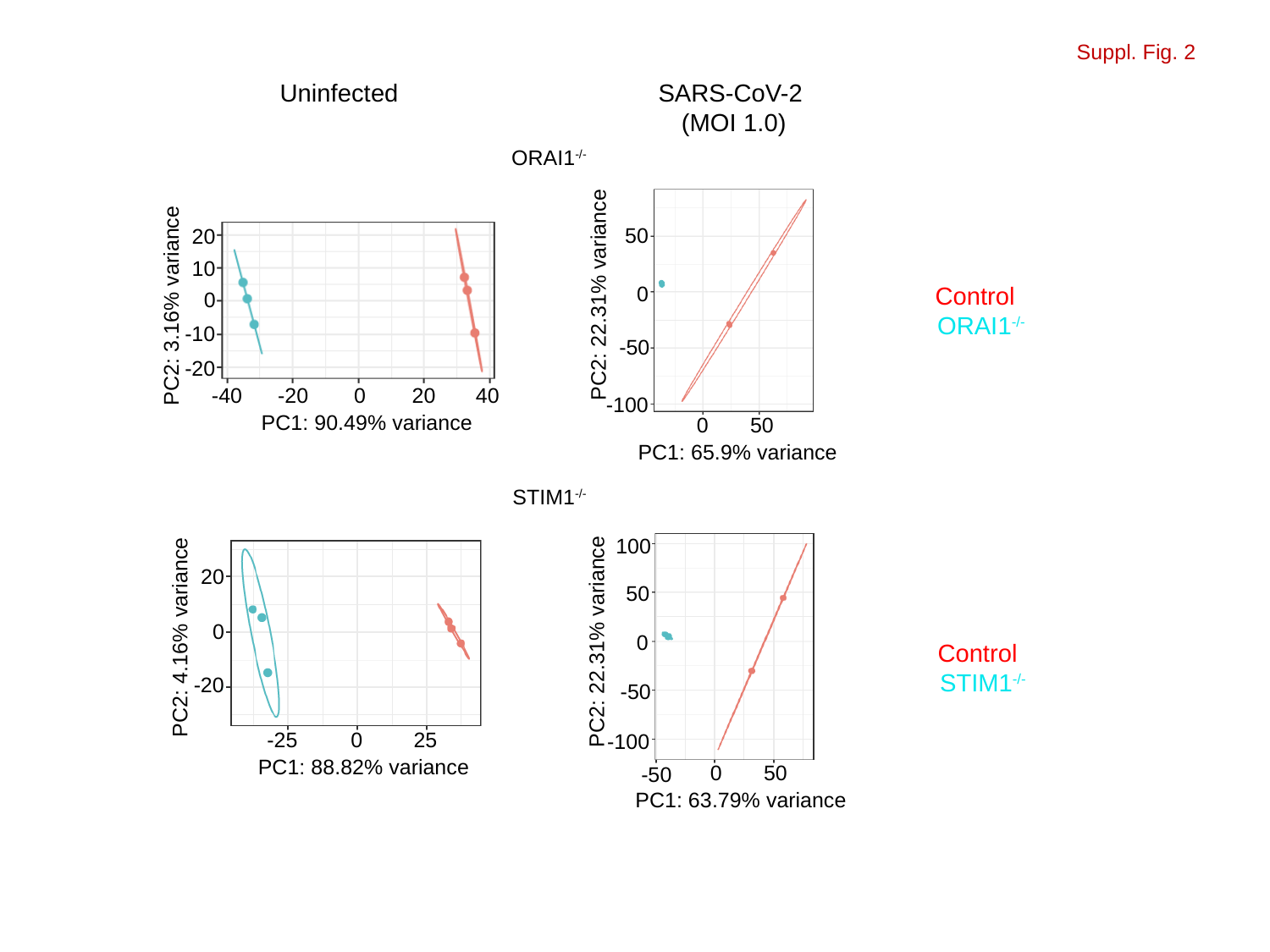

Suppl. Fig. 2
Uninfected
SARS-CoV-2 (MOI 1.0)
ORAI1^-/-^
50
20
10
0
Control
PC2: 22.31% variance
0
PC2: 3.16% variance
ORAI1^-/-^
-10
-50
-20
-40
-20
0
20
40
-100
PC1: 90.49% variance
0
50
PC1: 65.9% variance
STIM1^-/-^
100
20
50
0
PC2: 4.16% variance
PC2: 22.31% variance
0
Control
STIM1^-/-^
-20
-50
-25
0
25
-100
PC1: 88.82% variance
0
50
-50
PC1: 63.79% variance

### Slide 3
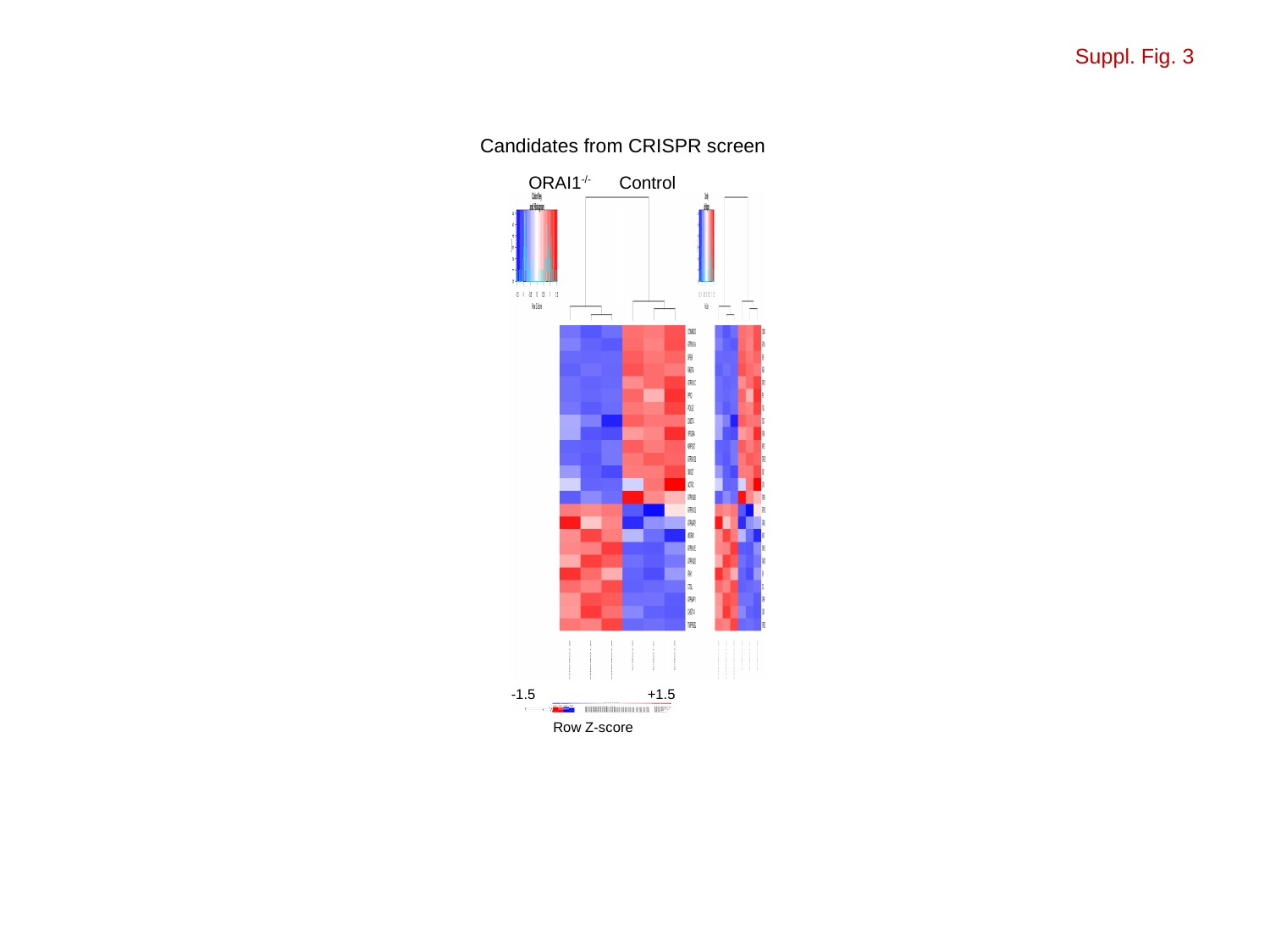

Suppl. Fig. 3
Candidates from CRISPR screen
ORAI1^-/-^
Control
-1.5
+1.5
Row Z-score
